# Establishing Design Principles for CRISPR-Cas Antifungals in *Candida albicans*

**DOI:** 10.64898/2026.08.27.747665

**Authors:** Christopher J. Cotter, Cong T. Trinh

## Abstract

Drug-resistant fungal pathogens pose a growing public health threat, causing millions of infections and deaths annually. Limited antifungal drug classes and rising resistance highlight the urgent need for novel therapies. CRISPR-Cas systems offer sequence-specific antimicrobial potential, but their efficacy is influenced by organism-specific DNA repair outcomes. Here, we demonstrate that in *Candida albicans*, which predominantly relies on homology-directed repair (HDR), both repair template availability and DNA repair enzyme activity critically determine Cas9-induced lethality. By providing Trojan Horse donor DNA repair templates when targeting essential and DNA repair genes, we show that Cas9 lethality can be selectively tuned. Furthermore, multiplexed gRNA targeting to modulate DNA repair capacity reveals strong synergistic interactions when co-targeting HDR components, which is corroborated by enhanced killing in HDR-compromised strains. These results establish DNA repair as a programmable determinant of CRISPR-Cas antifungal activity and provide a mechanistic framework for combinatorial targeting strategies, advancing the development of CRISPR-Cas antifungals.

**GRAPHICAL ABSTRACT:** 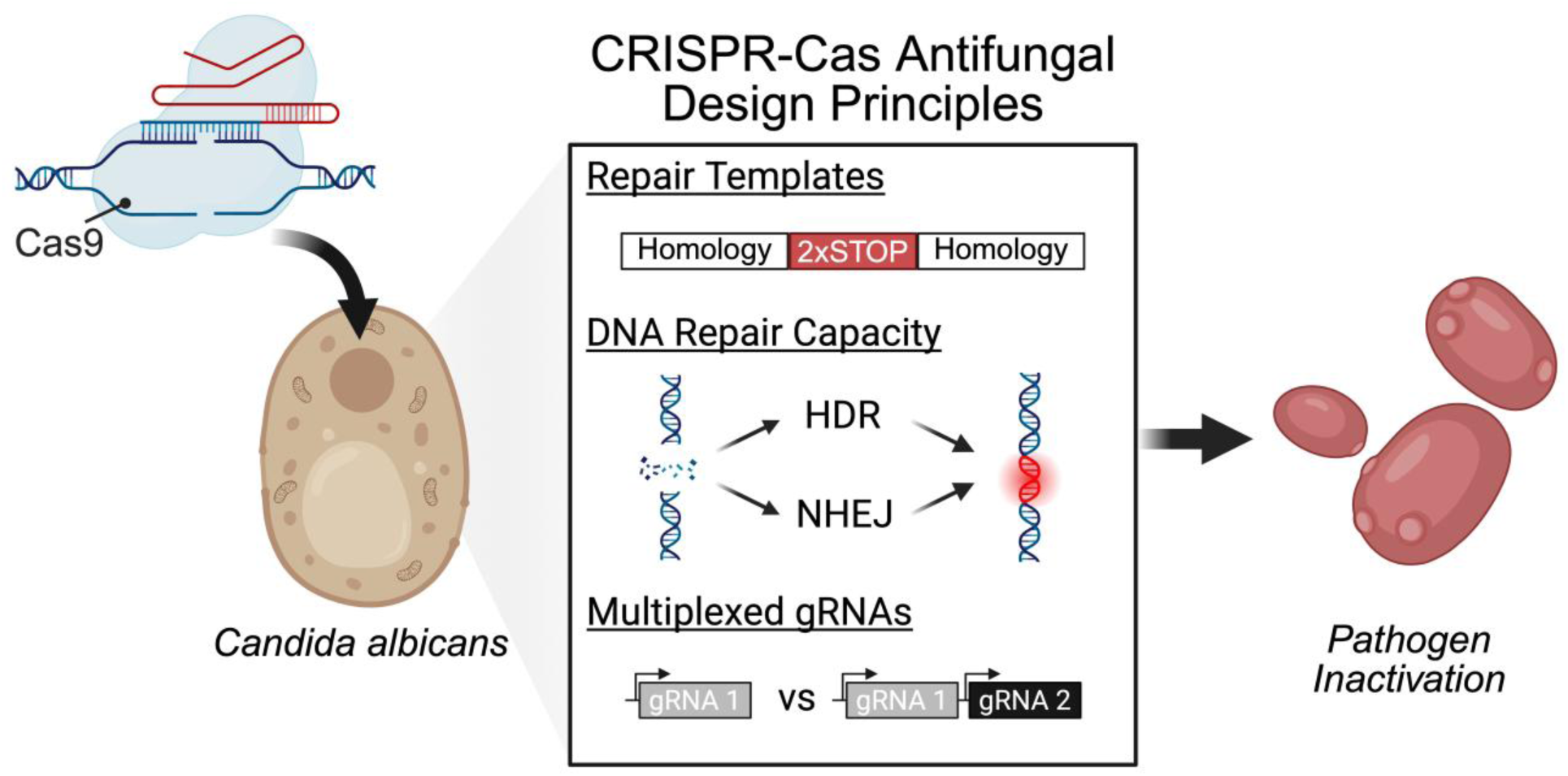

## INTRODUCTION

Drug-resistant fungal pathogens pose an immediate public health threat. Human fungal pathogens, including species from *Candida*, *Aspergillus*, and *Cryptococcus*, cause over one billion infections and an estimated 3.5 million deaths annually^1,2^. The prevalence of fungal infections is surging due to an increase in immunocompromised individuals, such as those undergoing chemotherapy or living with HIV/AIDS, as well as an increase in geographic range and dispersal due to climate change^3,4^. Along with an increased at-risk population, the threat of fungal pathogens is amplified by the rapid emergence of antifungal resistance and limited classes of antifungal drugs^5^. Overuse of current antifungals in both medical and agricultural settings has escalated the emergence of antifungal-resistant species (e.g., *Candida auris*)^6–10^. The rapid emergence and high mortality rates of multidrug-resistant isolates present an urgent need for new strategies to combat fungal pathogens.

CRISPR-Cas systems offer a promising approach to combat resistant and rapidly evolving pathogens and have recently been deployed as sequence-specific antimicrobials in bacteria^11–15^. By utilizing organism-specific guide RNAs (gRNAs), CRISPR-Cas antimicrobials can selectively target and cleave genes of interest causing persistent DNA damage that results in lethal phenotypes, reduced virulence, or re-sensitization to existing antibiotics^16,17^. As antifungal agents, CRISPR-Cas systems have the potential to eliminate the pathogen of interest with high specificity, minimizing off-target toxicity to the host and disruption of the surrounding microbiome associated with current antifungal therapies^18–21^. However, investigation into the design principles for formulating effective CRISPR-Cas antifungals is still limited.

CRISPR-Cas antimicrobials in bacteria operate on the premise that Cas9-induced DNA damage is inherently lethal irrespective of the gene target^15,16^. Because most bacteria are haploid and lack robust DNA repair mechanisms, DNA lesions often persist resulting in cell death^22^. Translating this strategy into eukaryotic pathogens such as fungi, however, presents a major technical hurdle. Fungi possess highly efficient and sophisticated DNA repair machinery, utilizing both homology-directed repair (HDR) and non-homologous end joining (NHEJ), making them highly resistant to double strand breaks^23,24^. Consequently, achieving potent CRISPR-Cas antifungals may require inhibiting DNA repair pathways and actively targeting genes essential for cell survival^22,25^.

The balance between HDR and NHEJ is species-dependent and shapes the outcome of CRISPR-induced damage. In the model yeast *Saccharomyces cerevisiae*, which is HDR-dominant but still utilizes NHEJ, our group observed that simultaneously co-targeting essential and DNA repair genes was more lethal than targeting either alone. Co-targeting essential genes with *RAD52*, a critical component of HDR, produced the strongest synergistic increase in Cas9-induced killing^25^. This synergy arises from NHEJ-mediated mutagenic repair of the *RAD52* locus, functionally disabling HDR and forcing Cas9-induced breaks in essential genes to either remain unresolved or be repaired by error-prone NHEJ, thereby amplifying lethal outcomes.

In contrast, *Candida albicans,* the most common cause of invasive candidiasis and a valuable model fungal pathogen, is diploid and rarely resorts to utilizing NHEJ^26–29^. Instead, *C. albicans* relies primarily on HDR to repair DSBs, rendering repair templates a key driver of Cas9-induced outcomes^24^. Repair templates may arise endogenously from homologous DNA, such as sister chromatids or replicated genomic regions, the abundance of which can vary with ploidy and genome replication dynamics. Alternatively, repair templates may be introduced exogenously, as is common in conventional CRISPR-mediated genome editing^30^. In HDR-dominant systems such as *C. albicans*, these templates can either suppress Cas9 lethality by mutating the target site and alleviating chromosomal injury or enhance killing by enforcing deleterious repair outcomes, depending on the function of the targeted gene. Currently, the use and control of repair templates for CRISPR-Cas antifungals in *C. albicans* remains poorly characterized.

In this study, we investigate DNA repair as a central and programmable determinant of CRISPR-Cas-mediated killing in the diploid pathogen *C. albicans*. By co-targeting essential and DNA repair genes, we demonstrate that Cas9 lethality can be deliberately tuned through repair pathway disruption and repair template-dependent outcomes. Multiplexed gRNA targeting reveals strong synergistic interactions between essential and DNA repair genes, establishing combinatorial targeting as a promising antifungal strategy. To further validate this model, we demonstrate that the lethality of single targeting gRNAs is enhanced in HDR-compromised backgrounds. Together, these findings establish co-targeting of essential and DNA repair genes as a generalizable CRISPR-Cas antifungal framework and provide a mechanistic blueprint for extending sequence-specific antimicrobials to clinically relevant fungal pathogens.

## MATERIALS AND METHODS

### Strains and Culturing Conditions

*C. albicans* SC5314 (ATCC MYA-2876) was used as the base strain for all experiments. All other strain information is listed in Table S1. *E. coli* NEB DH10β was used for cloning and cultured in LB media at 37°C. All *C. albicans* strains were routinely cultured in YPD media at 30°C. YPD plates supplemented with nourseothricin (GoldBio) at 250 µg/mL were used for selection of transformants.

### Single and Multiplexed gRNA Integrative Plasmid Construction

All plasmids, primers and gRNAs are listed in Tables S2-4 respectively. pV1093-CatRNA was used as the base plasmid for all integrative constructs. Single and multiplexed CRISPR-GRIT gRNAs were constructed using Golden Gate Assembly, as described in Cotter and Trinh^31^. Before transformation, 5-10 µg of DNA was linearized using KpnI and SacI in an overnight restriction digest at 37°C. The digests were purified using the Omega Biotek E.Z.N.A. Cycle Pure Kit and the total purified products were used for subsequent transformations.

### Autonomously Replicating Linear Plasmid Construction

Origin of replication sequences were PCR amplified from *C. albicans* SC5314 genomic DNA using primers CC-932/CC-933 (ORI410) and CC-934/CC-935 (ORI1046), and cloned into the NotI-digested pV1093-CatRNA backbone via Gibson assembly^32^. Prior to transformation, the ORI containing plasmids were linearized by PCR using primers CC-936 and CC-937 to append 1.5x telomere repeats (CCGTACACCAAGAAGTTAGACATCCGTACACCAA) to the 5’ and 3’ end respectively. PCR products were purified using the Omega Biotek E.Z.N.A. Cycle Pure Kit prior to transformation into *C. albicans*.

### Construction of DNA Repair Mutant Strains

DNA repair genes were disrupted using transient CRISPR-GRIT gRNAs (Table S2). *C. albicans* SC5314 was transformed with 1 µg of PCR linearized pCC-CaCas9-ORI1046 harboring *RAD51*, *RAD52*, or *LIG4* CRISPR-GRIT gRNAs. Colony PCR validated colonies were restreaked on YPD plates supplemented with 250 µg/mL nourseothricin to isolate homogeneous populations. Following isolation, biallelic gene disruptions were confirmed by Sanger sequencing of the target locus followed by TIDE analysis^33^. Linear plasmids were cured from cells by streaking cells on YPD plates lacking nourseothricin and incubating for 2 days at 30°C.

### High Efficiency Transformation of C. albicans

Transformations were carried out using the FACT method^34^. Briefly, single freshly streaked *C. albicans* colonies were used to inoculate in 10 mL of minimal media (6.7 g/L yeast nitrogen base without amino acids, 5 g/L glucose, and 0.36 g/L potassium acetate) and incubated for 12-16 h in a shaking incubator at 30°C, 250 rpm. Cells were diluted to an OD600 of 0.3 in 25 mL of minimal media and incubated at 30°C, 250 rpm for 5-6 h until the culture reached mid-exponential phase (OD600 of ∼1.8). Cells were collected by centrifugation at 900 × g for 5 min, washed once with 12 mL of sterile water, and resuspended to an OD600 of 30 in 100 mM lithium acetate titrated to a pH of 5 using glacial acetic acid. One hundred microliters of cells were aliquoted in sterile 1.5 mL tubes. One microgram of linearized DNA, 10 μL of 10 mg/mL herring sperm DNA, and 600 μL of 50% polyethylene glycol (PEG) 3350 dissolved in 100 mM lithium acetate were then added to each tube. Transformation mixtures were incubated for 18-20 h static at 30°C then heat shocked in a water bath at 44°C for 15 min. Cells were then pelleted at 900 × g for 5 min and washed with 1 mL of YPD. Cells were resuspended in 2 mL of YPD in a 15 mL culture tube and incubated for 4 h in a shaking incubator at 30°C and 250 rpm before plating serial dilutions on YPD agar plates supplemented with 250 μg/mL nourseothricin for selection. Plates were incubated at 30°C for 2 days before counting colonies.

### Calculation of Log₂ Colony Reductions and Interaction Scores

Following transformations, colony forming units (CFU) from each plate were counted using ImageJ^35^. Log_2_ colony reductions (log_2_CR) of targeting gRNAs relative to the non-targeting control were calculated as follows:

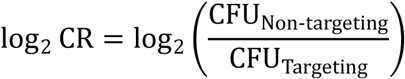

For multiplexed constructs, interaction scores were calculated from log_2_ colony reduction values using an adapted gene co-targeting interaction analysis formula^25^. Assuming gRNA co-targeting is additive and accounting for expression limitations in our multiplexed gRNA design, expected multiplexed colony reductions were calculated using the following equation:

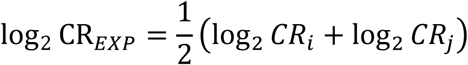

where i and j represent the individual gRNA targets in the multiplexed pair. The interaction score (IS) was then calculated as follows:

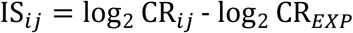

where log_2_CR_ij_ is the multiplexed log_2_CR for gRNAs i and j. Here a positive interaction score indicates synergy as the combination produced a greater than additive effect, while negative scores indicate antagonism.

### Statistical Analysis

Statistical analyses were performed using GraphPad Prism 10. Statistical significance was determined using unpaired two-tailed t-tests for comparisons between two groups, one-way ANOVA for single-factor analyses, and two-way ANOVA for experiments involving two independent variables, followed by Fisher’s LSD or Tukey’s post hoc tests where appropriate. Sample numbers, p-values, and the statistical test used for each analysis can be found in the respective figure legends.

## RESULTS

### Guide RNAs with Integrated Repair Templates as a Trojan Horse for Controlling the Potency of CRISPR-Cas Antifungals in *C. albicans*

DNA repair is a critical determinant of CRISPR-Cas antimicrobial outcomes. In HDR-dominant pathogens, cell survival following DNA damage depends on both the activity of DNA repair enzymes and the availability of repair templates. We hypothesize that modulating DNA repair by perturbing enzyme and repair template availability can enhance genome instability and maximize CRISPR-based antifungal activity.

To test this hypothesis, we first investigated the effect of repair template availability on Cas9-induced lethality in *C. albicans*. To eliminate variables associated with repair template delivery, we employed our CRISPR-GRIT (Guide RNAs with Integrated Repair Templates) system^31^. This system tethers a donor DNA repair template containing premature stop codons for gene disruption to the gRNA seed sequence, ensuring gRNA-repair template co-localization and a homogeneous distribution of repair templates across cells in a transformed population. This CRISPR-GRIT design serves as a “Trojan Horse” to exploit *C. albicans’* strong HDR machinery. To survive the Cas9-induced chromosomal damage, cells must incorporate the provided repair templates to eliminate the PAM sequence and prevent further cleavage. However, because the repair templates encode for premature stop codons, the cells inadvertently disrupt critical gene functions, trapping the cells and inducing lethal outcomes (Figure 1A). As a proof of concept for this system, we chose to evaluate its lethality against *C. albicans* homologs of top-performing essential (*RPS15, TRL1, RPB2*) and DNA repair genes (*RAD51, RAD52, LIG4*) identified in our *S. cerevisiae* study^25^ (Figure 1B).

**Figure 1.**
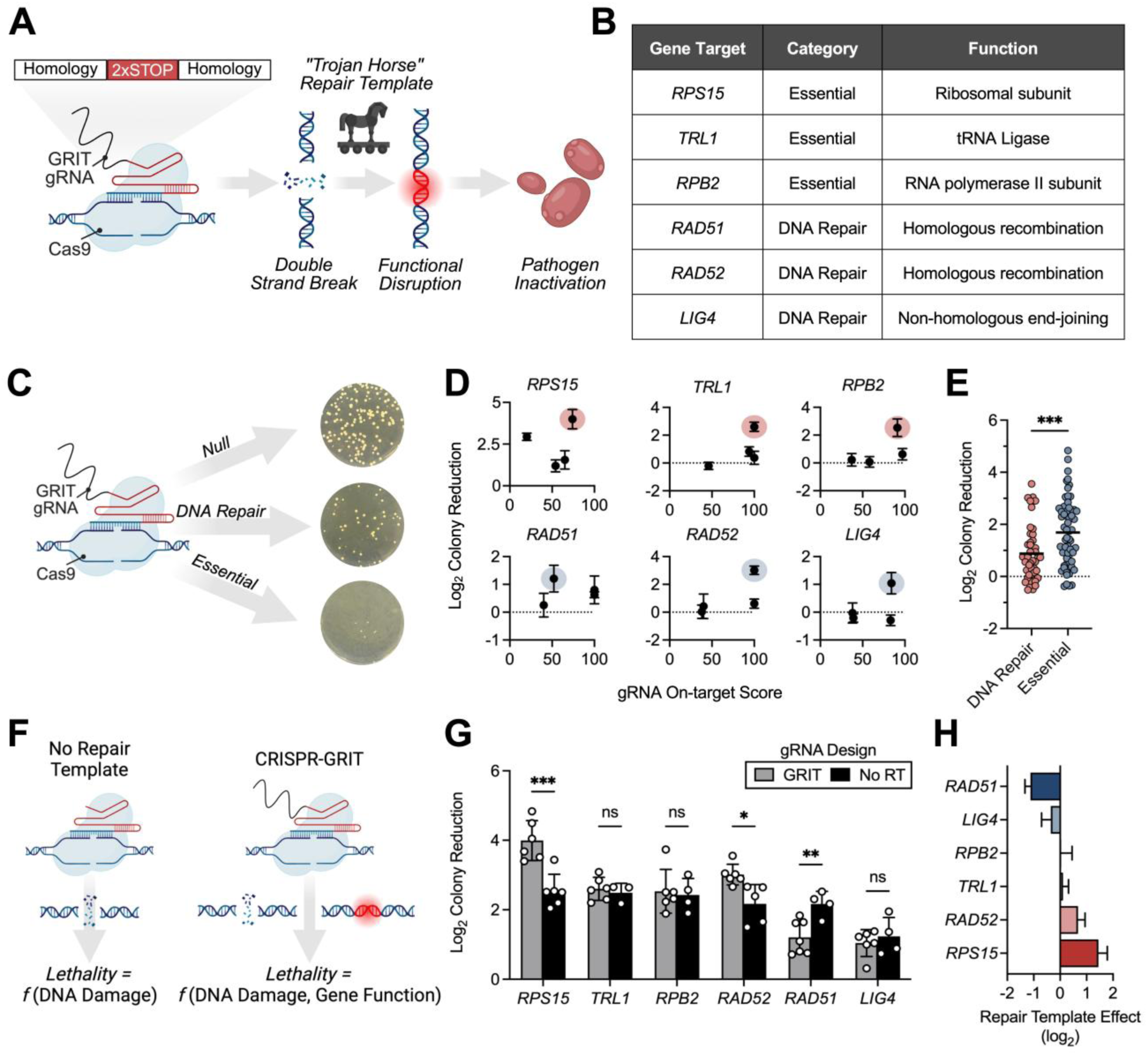
Repair Templates Differentially Modulate CRISPR-Cas Lethality in *C. albicans*. **(A)** Schematic of CRISPR-Cas lethality screening in *C. albicans*. **(B)** Identification of most lethal CRISPR-GRIT gRNAs for each gene target. (**C)** Comparison of lethality between essential and DNA repair targets. **(D)** Differences in modes of lethality between conventional CRISPR designs without repair templates and the CRISPR-GRIT design. **(E)** Colony reductions of conventional (No RT) and CRISPR-GRIT designs for the top gRNAs for each target. Black lines represent the mean of each dataset. **(F)** Difference in colony reductions between the CRISPR-GRIT design and the conventional design for each target. Data in **(B)**, **(C), (E)** are the mean ± standard deviation of at least three biological replicates. Data in **(D)** are the mean ± standard error of the mean of at least three biological replicates. Statistical significance for (**C)** and **(E)** was evaluated using unpaired t-tests. *P < 0.05, **P < 0.01, ***P < 0.001, ****P < 0.0001. Panels **(A), (C)**, and **(F)** were created in BioRender. Trojan horse icon in Panel **(A)** is modified from “Trojan horse” by Freepik (Magnific), on Flaticon (https://www.flaticon.com/free-icon/trojan-horse_5830744).

### Potency of CRISPR-Cas Antifungals Depends on Targeted Gene Functions

Using our CRISPR-GRIT system, we first screened for highly active gRNAs in a transformation-based lethality assay. Here, single-targeting Cas9-gRNA constructs were transformed into *C. albicans* and transformation efficiencies of each construct were compared to a non-targeting control (Figure 1C). For each target gene, we tested four CRISPR-GRIT gRNAs with different predicted on-target scores to account for gRNA-specific variability and identify the most effective guides (Figure 1D, Table S5). Highly active gRNAs are expected to produce more efficient Cas9-mediated cleavage, resulting in extensive DNA damage which reduces cell viability and creates selection pressure to incorporate the Trojan Horse repair templates.

Transformation of these single-target CRISPR-GRIT constructs resulted in variable reductions in colony survival, with differences observed both between target genes and among individual gRNAs for the same target (Figure 1D, Table S5). The most active guides for each gene, CaVip-3 (*RPS15*), CaVip-7 (*TRL1*), CaVip-12 (*RPB2*), CaVip-15 (*RAD52*), CaVip-17 (*RAD5*1), and CaVip-23 (*LIG4*), were chosen for use in subsequent experiments. The observed variability among gRNAs reflects differences in Cas9-gRNA cleavage efficiency as well as target-dependent sensitivity to CRISPR-mediated gene disruption. As expected, gRNAs targeting essential genes were on average more lethal than those targeting DNA repair genes due to the direct disruption of cellular processes required for cell survival (Figure 1E).

Among the essential genes, *RPS15* was the most lethal target, resulting in a 16-fold decrease in viable colonies relative to the non-targeting control (Figure 1D, Table S5). *RPS15*, a putative ribosomal protein, has been reported to be haploinsufficient under certain conditions, where the loss of a single allele can reduce viability^36^. Because single-copy repair template integration is kinetically favored over biallelic disruption, this vulnerability amplifies lethality using the Trojan Horse design.

In contrast, loss of DNA repair genes *RAD51* and *LIG4* was generally tolerated (Figure 1D, Table S5). Disruption of *RAD52*, however, was unexpectedly lethal, with the top-performing gRNA causing 8-fold colony reduction relative to the non-targeting control, comparable to that observed for essential gene targets (Figure 1D, Table S5). Although primarily involved in homologous recombination, *RAD52* also plays a role in maintaining chromosome stability, the disruption of which can cause genotoxic stress and growth defects.

### Repair Templates Differentially Modulate the Potency of CRISPR-Cas Antifungals

To better understand how our CRISPR-GRIT design impacts Cas9-induced lethality, we compared it to conventional gRNA designs containing 20 bp spacers without repair templates using the top-performing gRNAs for each gene target (Figure 1F). Across the six targets, we observed target-dependent differences in lethality between the two designs (Figure 1G-H). In the absence of repair templates (conventional gRNAs), lethality was similar across all targets, indicating comparable gRNA activity and a relatively constant contribution of DNA damage to lethality irrespective of the target locus (Figure 1G). However, the introduction of repair templates via the CRISPR-GRIT design resulted in divergent target-dependent outcomes.

For essential genes, the *RPS15* gRNA exhibited a 1.6-fold increase in lethality (p < 0.01) with the CRISPR-GRIT design relative to the conventional gRNA, whereas *TRL1* and *RPB2* gRNAs showed no increase (Figure 1G-H). For DNA repair targets, CRISPR-GRIT-mediated disruption of *RAD52* led to a slight increase in lethality, while disruption of *RAD51* resulted in reduced lethality, and *LIG4* showed no significant difference (Figure 1G-H). These results demonstrate that while CRISPR-induced DNA damage contributes a largely uniform lethal burden, repair template-mediated editing modulates cell survival in a bidirectional, target-dependent manner.

### Multiplexed CRISPR-Cas Antifungals Enhance Combinatorial Synergy

Next, we evaluated the effect of co-targeting essential and DNA repair genes on the potency of CRISPR-Cas antifungals in *C. albicans*. We designed multiplexed constructs combining top-performing CRISPR-GRIT gRNAs arranged in discrete transcriptional units and transformed them into *C. albicans* to determine their effect on cell viability (Figure 2A). To ensure that the position of the gRNA within the expression cassette did not bias activity, we tested both possible orientations (e.g., gRNA1-gRNA2 and gRNA2-gRNA1). We observed no significant orientation-dependent effects on lethality (Figure S1), allowing us to average the reciprocal data for each gRNA pair. On average, multiplexed gRNAs produced greater lethality than the average single-target construct (Figure 2B), with combinations containing either *RPS15* or *RAD52* gRNAs consistently exhibiting the highest overall colony reduction (Figure 2C).

**Figure 2.**
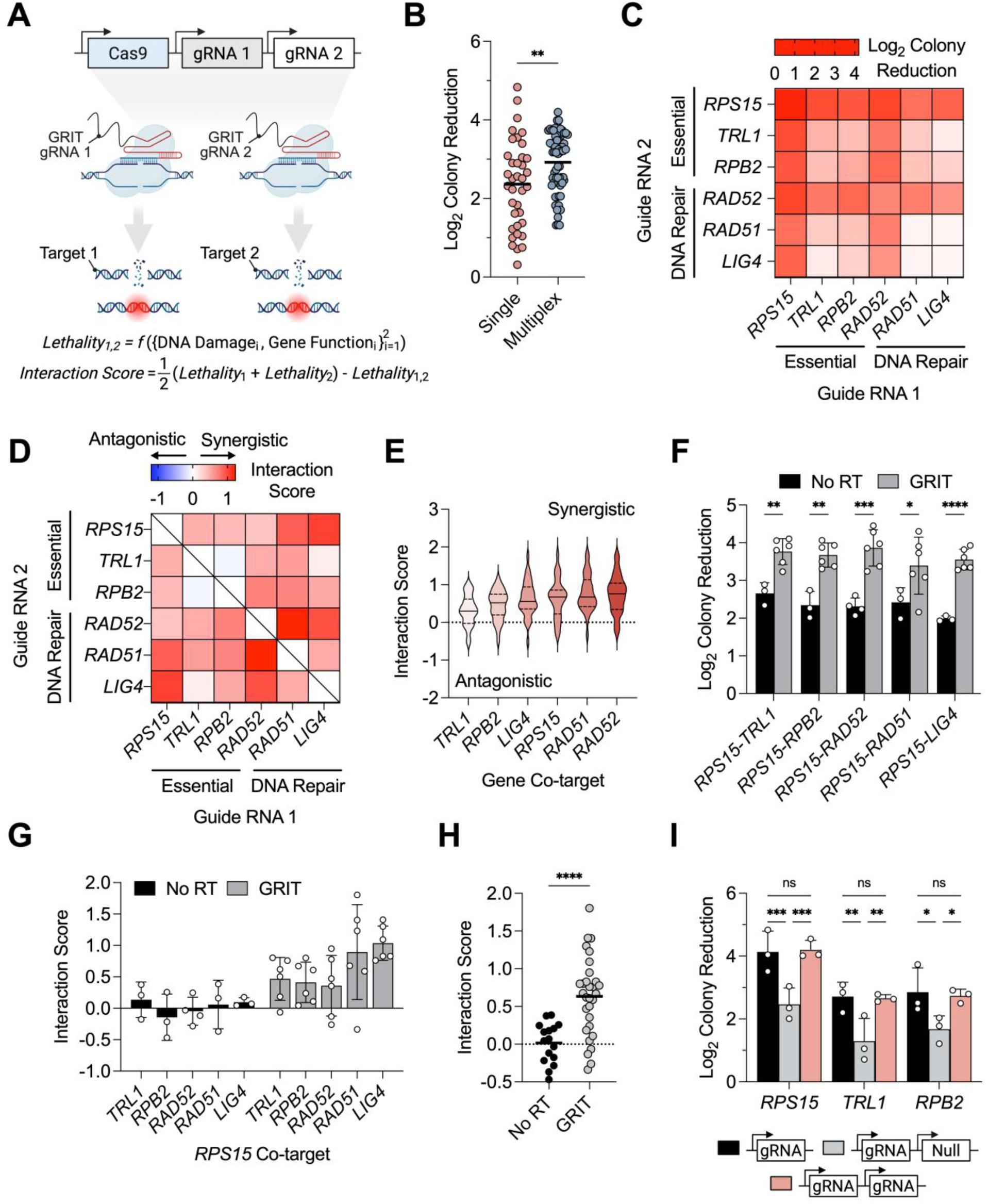
Multiplexed CRISPR-GRIT gRNAs Produce Synergistic Lethality in *C. albicans*. **(A)** Schematic of multiplexed CRISPR-Cas targeting and interaction scoring. **(B)** Comparison of single and multiplexed lethality across all targets. (**C)** Heat map of colony reductions for each multiplexed target pair. **(D)** Heat map of interaction scores for each multiplexed target pair. Self-paired gRNA constructs (e.g., *RPS15* + *RPS15*) serve as technical controls rather than true combinatorial gRNA interactions and were therefore excluded from the interaction score heatmap as indicated by the crossed-out diagonal. White off-diagonal cells denote an interaction score of zero (additive effect). **(E)** All interaction scores for pairs containing a particular gene co-target. **(F)** The effect of CRISPR-GRIT repair templates on multiplexed colony reductions for a subset of combinations containing the *RPS15* gRNA. **(G)** Interaction scores with and without integrated repair templates for combinations with *RPS15.* **(H)** Comparison of interaction scores across designs with and without repair templates for the *RPS15* multiplexed subset. **(I)** Colony reductions of essential CRISPR-GRIT gRNAs in single targeting designs or multiplexed with either a null gRNA or itself. Data are the mean ± standard deviation of at least three biological replicates. Statistical significance was evaluated using unpaired t-tests **(B)** and two-way ANOVA **(F)**. *P < 0.05, **P < 0.01, ***P < 0.001, ****P < 0.0001. For dot plots in **(B)** and **(F)**, black lines represent the mean of each dataset. Panel **(A)** was created in BioRender.

To quantitatively assess gRNA interactions, we applied a modified gene co-targeting interaction model^25^. Here, interaction scores greater than 0 indicate synergy, scores less than 0 suggest antagonism, and scores equal to 0 indicate that the gRNAs are purely additive. This analysis revealed that multiplexed pairings were broadly synergistic, with the exception of *TRL1* paired with *RPB2* (Figure 2D). Co-targeting any gene with *RAD52* produced particularly strong synergy suggesting that compromising HDR can enhance CRISPR-Cas lethality (Figure 2E). The combination of *RAD51* and *RAD52* exhibited the strongest synergy (mean interaction score = 1.23). Since disruption of either gene compromises HDR, targeting both amplifies the lethal effect of the CRISPR-induced DNA damage.

### Repair Templates Drive CRISPR-Cas Antifungal Synergy

To determine how repair templates impact multiplexed gRNA synergy, we compared lethality and interaction scores across a subset of *RPS15*-containing combinations with and without the integrated repair templates. Consistent with our single-target observations, the CRISPR-GRIT design was consistently more lethal than conventional multiplexed gRNAs lacking repair templates (Figure 2F). Moreover, when evaluating interaction scores for the *RPS15* subset, all combinations utilizing the CRISPR-GRIT design were synergistic (mean interaction score of 0.63), whereas the corresponding conventional gRNA combinations lacking repair templates were simply additive (mean interaction score of 0.017; Figure 2G-H). These findings demonstrate that repair templates not only modulate lethality but fundamentally dictate the nature of the gRNA interactions by enabling functional disruption of target genes.

### Multiplexed Lethality is Impaired by Expression Limitations

Despite the strong synergistic interactions, we observed that the top-performing single target constructs displayed higher absolute lethality than the top multiplexed constructs (Figure 2B). To investigate this limitation, we multiplexed the top-performing essential CRISPR-GRIT gRNAs either with a null-targeting gRNA or with themselves (Figure 2I). Multiplexing with a null-targeting gRNA reduced lethality by approximately two-fold across all tested guides compared to single-gRNA expression (Figure 2I). In contrast, pairing each gRNA with itself restored lethality to levels comparable to the single-gRNA system (Figure 2I). These findings indicate that introducing additional gRNAs dilutes Cas9 activity among competing guides, suggesting that Cas9 availability or gRNA expression is limiting in our system. Therefore, while multiplexing successfully revealed synergistic vulnerabilities, further optimization of Cas9 and gRNA expression will be required to maximize the therapeutic potency of combinatorial targeting.

### CRISPR-Cas Lethality is Enhanced in Homologous Recombination Deficient Strains

Since DNA repair functional knockouts appear to be critical for co-targeting synergy, we hypothesized that CRISPR-induced lethality would increase in DNA repair gene knockout strains. To test this, we constructed Δ*RAD51,* Δ*RAD52,* and Δ*LIG4* background strains and transformed them with our single targeting CRISPR-GRIT constructs. Since *RAD51* and *RAD52* are required for homologous recombination, the integrative Cas9 system used in the initial experiments failed to transform efficiently into these deletion strains, limiting our dynamic range for evaluating Cas9-induced lethality (Figure 3C).

**Figure 3.**
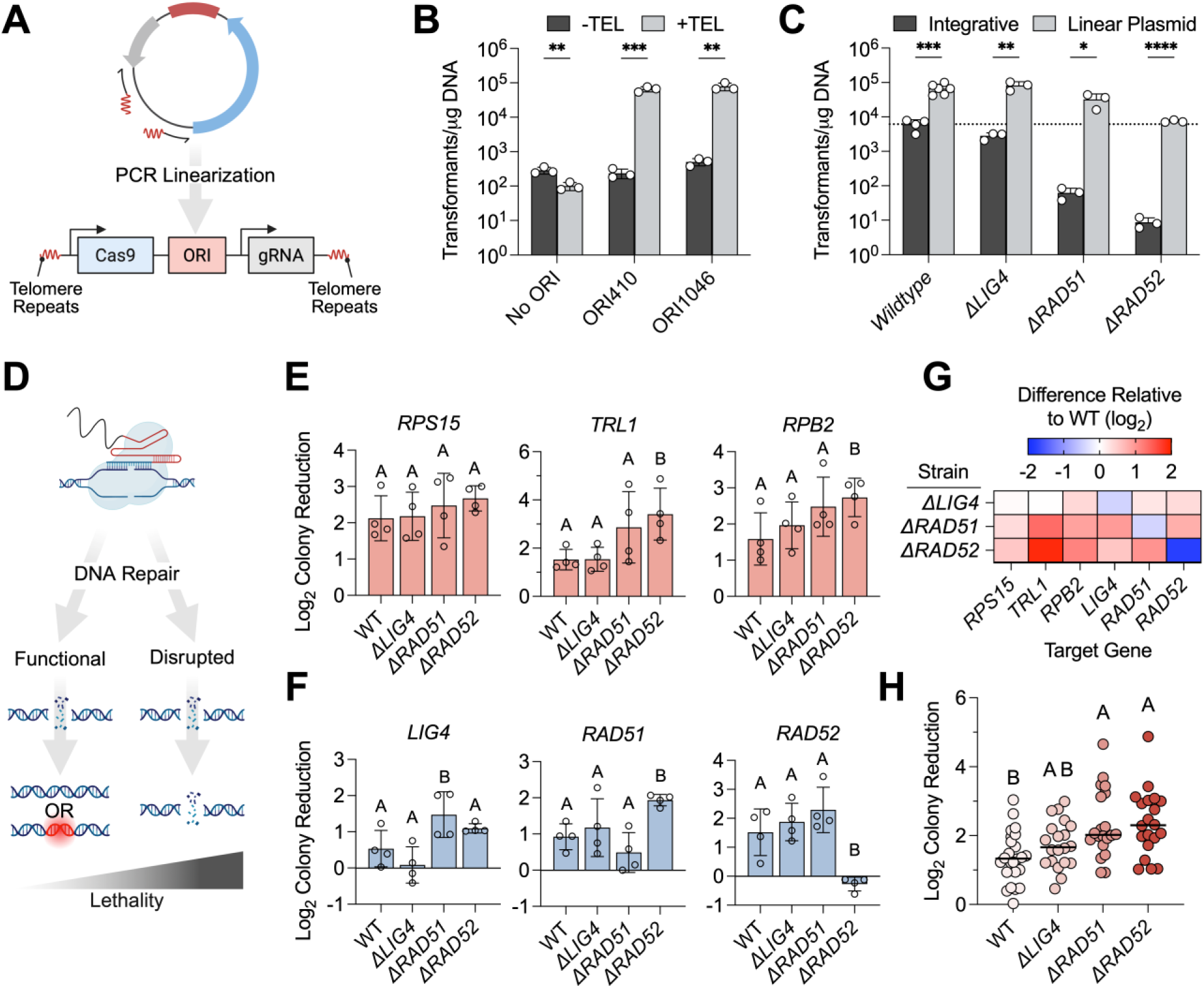
CRISPR Lethality is Enhanced in DNA Repair Deletion Strains. **(A)** Autonomously replicating linear CRISPR-Cas plasmid design. **(B)** Transformation efficiencies of linear plasmids containing different ORI sequences with and without 1.5x telomere repeats capping the DNA ends. (**C)** Comparison of transformation efficiencies using linear plasmids harboring ORI1046 and 1.5x telomere repeats against integrative CRISPR-Cas constructs in DNA repair deletion strains. Statistical significance for **B** and **C** was evaluated using unpaired t-tests. **(D)** Schematic of lethality outcomes in cells with functional or disrupted DNA repair genes. **(E-F)** Colony reductions of linear plasmids harboring CRISPR-GRIT gRNAs targeting **(E)** essential (*RPS15, TRL1, RPB2*) and **(F)** DNA repair (*LIG4, RAD51, RAD52*) genes in WT and DNA repair deletion strains (*ΔLIG4, ΔRAD51, ΔRAD52*). Statistical significance was assessed using one-way ANOVA followed by Fisher’s LSD post-hoc test comparing each DNA repair deletion strain to the WT strain. Groups labeled A are not significantly different from the WT, whereas groups labeled B are significantly different from the WT (P < 0.05). **(G)** Heat map of change in colony reduction in DNA repair deletion strains relative to the WT strain. **(H)** Comparison of CRISPR-Cas antifungal lethality across different DNA repair deletion strains. Black lines represent the median of each dataset. Statistical significance was determined using one-way ANOVA followed by Tukey’s post-hoc test. Statistical significance is denoted with compact letter display where groups with different letters are statistically significant from one another (P < 0.05) and groups with the same letter are not. All data are the mean ± standard deviation of at least three biological replicates. *P < 0.05, **P < 0.01, ***P < 0.001, ****P < 0.0001. Panels **(A)** and **(D)** were created in BioRender.

To overcome this, we constructed a linear CRISPR-Cas plasmid capable of autonomous replication without chromosomal integration, allowing it to transform efficiently in both wildtype and HDR-deficient strains (Figure 3A-C). Testing two previously characterized origins of replication (ORI410 and ORI1046)^37^ with and without PCR-added telomere repeats revealed that telomere repeats coupled with a bona fide ORI were critical, increasing the transformation efficiency by over 150-fold for constructs with either ORI (Figure 3B). For subsequent experiments, we proceeded with ORI1046, which transformed efficiently in both wildtype and DNA repair deletion strains (Figure 3C). We validated that the CRISPR-Cas linear plasmid system was functional by targeting both the *ADE2* and *ACE2* genes with our previously utilized CRISPR-GRIT gRNAs, where homozygous deletions resulted in either red or wrinkled colonies, respectively (Figure S2).

Using these linear Cas9 plasmids, we then transformed wildtype and DNA repair deletion strains (*ΔRAD51*, *ΔRAD52*, and *ΔLIG4*) with single-target CRISPR-GRIT constructs to evaluate the impact of compromising DNA repair pathways on Cas9-induced lethality (Figure 3D). Consistent with our hypothesis, disruption of HDR sensitized cells to Cas9-induced DNA damage. Assessing lethality across our panel of essential (Figure 3E) and DNA repair (Figure 3F) targets revealed that the *ΔRAD52* background strain yielded statistically significant increases in lethality specifically when targeting *TRL1*, *RPB2*, and *RAD51*. To visualize the broader landscape of this sensitization, we mapped the difference in colony reduction for each target in the knockout strains relative to the wildtype baseline (Figure 3G). Despite the observed target-to-target variability, aggregate analysis of all tested guides demonstrated that strains lacking either *RAD51* or *RAD52* exhibited significantly greater Cas9-induced colony reductions (median log_2_ colony reductions of 2.02 and 2.30 respectively, p < 0.001) compared to the wildtype strain (median 1.32; Figure 3H). In contrast, the *ΔLIG4* strain showed no significant difference in lethality, reflecting its limited role in *C. albicans* DNA repair (Figure 3E-F). These findings reinforce the synergy observed when co-targeting *RAD51* and *RAD52* and confirm that HDR is the dominant repair pathway in *C. albicans*. By eliminating HDR capacity, the lethal outcomes of Cas9-induced DNA damage can be amplified, validating DNA repair as a programmable determinant of CRISPR-Cas antifungal activity.

## DISCUSSION

Drug-resistant fungal pathogens represent a growing public health challenge, and the limited number of antifungal drug classes constrains therapeutic options. CRISPR-Cas systems offer a versatile, programmable solution to combat fungal pathogens; however, the design rules for creating effective CRISPR-antifungals are poorly understood. In this study, we define DNA repair capacity as a central and programmable driver of CRISPR-Cas-mediated killing in the diploid fungal pathogen *C. albicans*. By systematically modulating repair template availability and disrupting DNA repair pathways, we show that Cas9-induced lethality is not dictated solely by chromosomal injury, but instead reflects the balance between DNA damage and repair efficiency. These findings establish DNA repair as a tunable parameter that can be leveraged and exploited to control the efficacy of sequence-specific CRISPR-based antifungal strategies.

In HDR-dominant organisms such as *C. albicans*, the abundance of homologous repair templates strongly influences the outcome of Cas9-induced DSBs. Endogenous templates derived from homologous chromosomes or replicated genomic regions can suppress lethality by enabling efficient repair, thereby limiting the effectiveness of CRISPR-based killing. Here, we demonstrate that this repair bias can be deliberately manipulated by delivering Trojan Horse repair templates. The CRISPR-GRIT design exploits *C. albicans’* strong HDR preference by supplying donor templates that enforce deleterious repair outcomes through the introduction of premature stop codons and PAM-disrupting mutations. In this context, the supplied repair templates not only facilitate genome editing but directly shape the balance between survival and lethality following target cleavage.

Our results show that the effect of repair templates is largely governed by the target gene’s function and dosage sensitivity. Among the essential genes, only *RPS15*, a haploinsufficient target, demonstrated a significant increase when a repair template was present. This observation suggests that dosage-sensitive genes may represent a strategic vulnerability for enhancing CRISPR antifungal efficacy in diploid or polyploid pathogens where monoallelic disruption is kinetically favored. DNA repair targets displayed divergent behaviors in the presence of repair templates with lethality increasing for *RAD52* but decreasing for *RAD51* and showing no change for *LIG4*. Disruption of *RAD52* not only compromises DNA repair capacity, but also induces broader chromosomal instability and fitness defects. In contrast, *RAD51* and *LIG4* mutations are generally tolerated and therefore template-directed repair alleviates Cas9-induced DNA damage at the target locus with no additional fitness costs.

Beyond influencing single-target outcomes, *C. albicans* HDR-proficiency also hinders CRISPR-Cas co-targeting strategies by suppressing loss-of-function mutations, an essential step for modulating DNA repair capacity and maximizing CRISPR antifungal potency. The CRISPR-GRIT design facilitates functional knockout of DNA repair enzymes themselves, enabling combinatorial targeting designs that compromise repair capacity while simultaneously inducing lethal DNA damage. This design feature is critical for interrogating, and exploiting, genetic interactions among gene pairs.

Consistent with this framework, multiplexed targeting revealed strong synergistic interactions when homologous recombination components were co-targeted, with *RAD51-RAD52* targeting constructs producing the greatest synergy. These findings closely parallel our prior work in *S. cerevisiae*, where co-targeting the essential HDR factor *RAD52* similarly amplified CRISPR-mediated killing^25^. Together, these studies indicate that repair pathway co-targeting represents a conserved strategy for enhancing CRISPR lethality across fungal species, despite differences in ploidy and DNA repair pathway usage. This cross-species consistency supports the conclusion that DNA repair capacity imposes a fundamental constraint on CRISPR antifungal efficacy.

More broadly, our results support a model in which CRISPR-mediated killing is governed by dominant DNA repair mode. In HDR-dominant pathogens, repair template availability plays a central role in determining CRISPR outcomes, rendering donor template design and availability a critical parameter for tuning lethality. In contrast, in NHEJ-dominant pathogens such as *Aspergillus* and *Cryptococcus* species, repair templates may contribute less directly to survival following DNA cleavage, limiting the applicability of CRISPR-GRIT-style strategies. Nevertheless, the underlying principle of co-targeting DNA repair pathways remains relevant. In such systems, disrupting key NHEJ components such as *KU70, KU80,* or *LIG4* may similarly sensitize cells to CRISPR-induced damage by reducing repair efficiency and increasing the persistence of lethal lesions^38,39^. Thus, while specific implementations must be adapted to the repair architecture of each pathogen, repair pathway disruption emerges as a generalizable strategy for enhancing CRISPR-based antifungals.

While this study focuses on defining genetic and mechanistic design principles for CRISPR-based antifungals, translational bottlenecks must be solved to realize the potential of these systems for therapeutic applications. Expression limitations as observed in this study must be addressed to enable efficient multiplexed editing and translate synergistic co-targeting interactions into superior absolute lethality. Additionally, fungal-specific nucleic delivery approaches, such as nanoparticles, conjugative elements^40^, or potentially mycovirus-derived platforms^41^, must be engineered for introducing CRISPR components into pathogenic fungi *in vivo*.

In summary, we demonstrate that DNA repair capacity and repair template availability are programmable determinants of CRISPR-Cas-mediated killing in *C. albicans.* By integrating repair template engineering with repair pathway co-targeting, this study establishes a framework for tuning CRISPR antifungal activity. These findings provide a mechanistic blueprint for extending sequence-specific antimicrobials to clinically relevant fungal pathogens and addressing the growing threat of antifungal resistance.

## ACKNOWLEDGEMENTS

Graphical abstracts and some conceptual figures 1A, 1C, 1F, 2A, 3A, and 3D were created in Biorender.

## Author contribution

Conceptualization (CTT), Investigation (CJC), Formal Analysis (CJC, CTT), Methodology (CJC, CTT), Validation (CJC), Visualization (CJC, CTT), Funding acquisition (CTT), Project Administration (CTT), Writing – original draft (CJC, CTT), Writing – review & editing (CJC, CTT).

## DATA AVAILABILITY

Supplementary Data are available online at NAR. Figure S1 demonstrates the effect of multiplexed gRNA orientation on CRISPR-Cas lethality. Figure S2 provides validation of the linear CRISPR-Cas plasmid system targeting *ADE2* and *ACE2*. All strains, plasmids, primers, and gRNAs used in this study are listed in Tables S1-4, respectively. Table S5 presents gRNA on-target scores and colony reduction values measured for each CRISPR-GRIT construct.

## CONFLICT OF INTEREST

CTT and CJC are named inventors on a U.S. Provisional Patent Application filed by the University of Tennessee Research Foundation (UTRF).

## FUNDING

This research is financially supported in part by the UT-ORII Seed Fund, the DoD ARL grant (W911NF261A160), the NSF grant (NSF2619053), and the NIH grant (1R21AI199008-01). The views, opinions, and/or findings contained in this article are those of the authors and should not be interpreted as representing the official views or policies, either expressed or implied, of the funding agencies. The mention of trade names or commercial products in this publication is solely for the purpose of providing specific information and does not imply a recommendation or endorsement by the funding agencies.

## Notes

### Competing Interest Statement

The authors have declared no competing interest.

