## Supplementary Figures S1-S2 and Tables S1-S5 for "Establishing Design Principles for CRISPR-Cas Antifungals in *Candida albicans*"

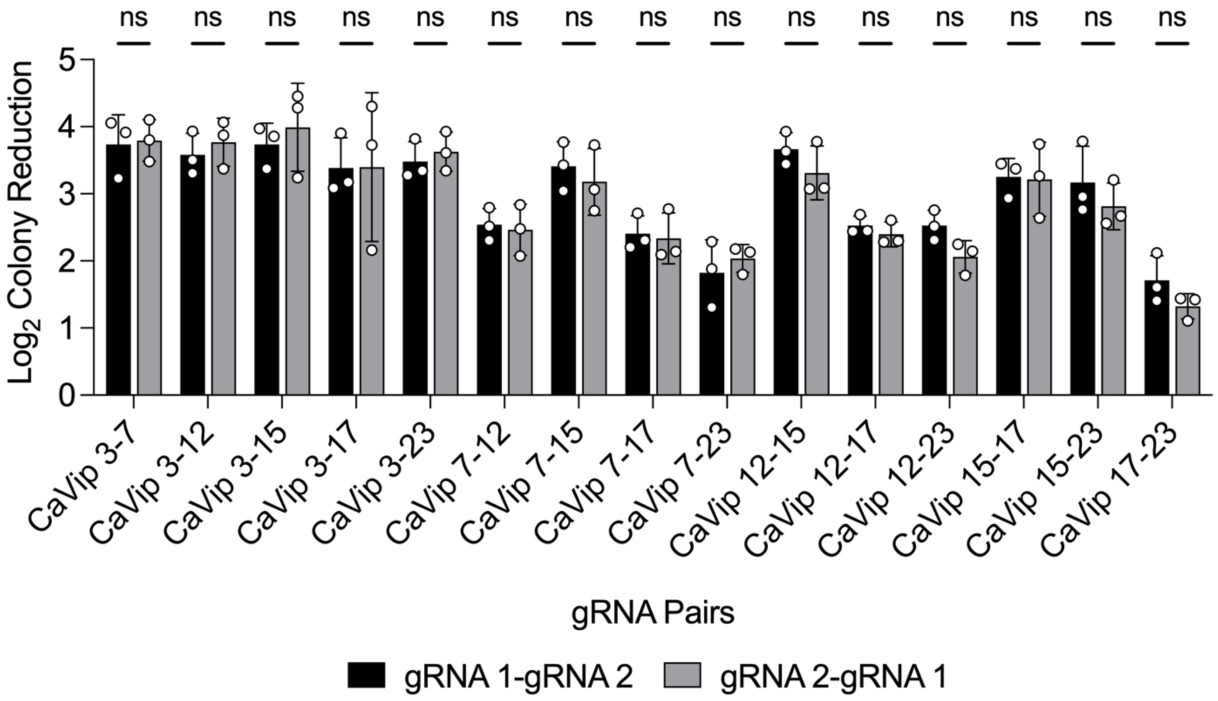


**Figure S1. Effect of gRNA orientation on CRISPR-Cas Lethality.** To determine if the position of the gRNA within the multiplexed expression cassette biased Cas9-induced lethality, reciprocal orientations were constructed and tested for all combinations. Each CRISPR-GRIT gRNA was arranged in a discrete transcriptional unit driven by its own *SNR52* promoter. X-axis labels represent the specific gRNAs paired in the construct (e.g., "CaVip 3-7" harbors both CaVip-3 and CaVip-7 gRNAs). Both possible orientations (e.g., p*SNR52*-CaVip-3::p*SNR52*-CaVip-7 and p*SNR52*-CaVip-7::p*SNR52*-CaVip-3) are represented as black and gray bars respectively. Data are the mean ± standard deviation of three biological replicates. Statistical significance was evaluated using multiple unpaired t-tests. No significant difference (ns) was observed between orientations for any construct ($p > 0.05$) demonstrating that gRNA position does not influence multiplexed lethality. Consequently, data from both orientations were pooled for the heatmap in Figure 2C and all subsequent interaction score analyses (Figures 2D-E).

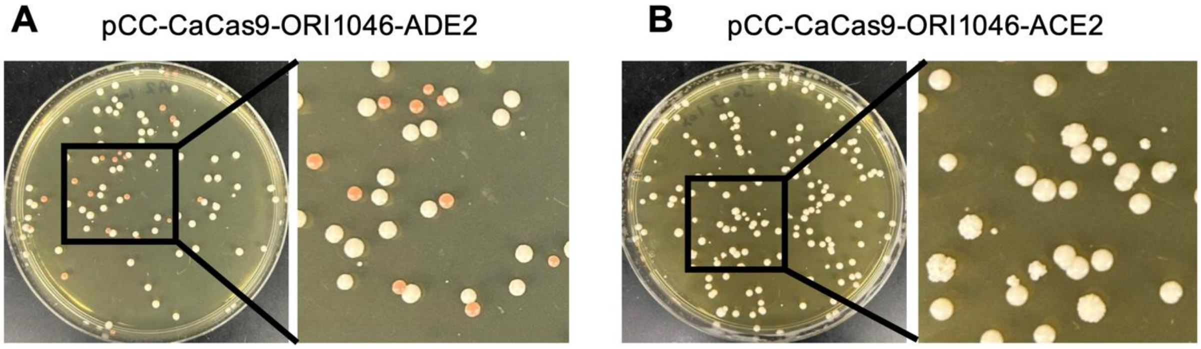


**Figure S2. Functional validation of CRISPR-Cas linear plasmid system. (A)** Plate pictures of *C. albicans* transformed with linearized pCC-CaCas9-ORI1046 harboring CRISPR-GRIT gRNAs targeting *ADE2* **(A)** where homozygous deletions result in red colony phenotypes and *ACE2* **(B)** with homozygous deletions resulting in wrinkled colony phenotypes.

**Table S1:** List of strains used in this study.

| Strain Name | Parental Strain | Genotype | Source |
|---|---|---|---|
| *C. albicans SC5314* | NA | Wildtype | ATCC (MYA-2876) |
| CaCC007 | *C. albicans SC5314* | *ΔRAD51* | This study |
| CaCC008 | *C. albicans SC5314* | *ΔRAD52* | This study |
| CaCC009 | *C. albicans SC5314* | *ΔLIG4* | This study |

| **Table S2:** List of plasmids used in this study. | | |
|---|---|---|
| **Plasmid** | **Description** | **Source** |
| pV1093-CatRNA | pV1093 based integrative CRISPR-Cas9 system with a tRNA-Ala at the 3' end of the SNR52 promoter for increased editing efficiency | Cotter & Trinh[1] |
| pV1093-Empty | pV1093-CatRNA based integrative CRISPR-Cas9 system with the SNR52 promoter and a BsmBI stuffer for gRNA multiplexing | Cotter & Trinh[1] |
| CaVip-1 | pV1093-CatRNA based integrative CRISPR-Cas9 system with a CRISPR-GRIT gRNA targeting RPS15 (Guide #1) | This study |
| CaVip-2 | pV1093-CatRNA based integrative CRISPR-Cas9 system with a CRISPR-GRIT gRNA targeting RPS15 (Guide #2) | This study |
| CaVip-3 | pV1093-CatRNA based integrative CRISPR-Cas9 system with a CRISPR-GRIT gRNA targeting RPS15 (Guide #3) | This study |
| CaVip-4 | pV1093-CatRNA based integrative CRISPR-Cas9 system with a CRISPR-GRIT gRNA targeting RPS15 (Guide #4) | This study |
| CaVip-5 | pV1093-CatRNA based integrative CRISPR-Cas9 system with a CRISPR-GRIT gRNA targeting TRL1 (Guide #1) | This study |
| CaVip-6 | pV1093-CatRNA based integrative CRISPR-Cas9 system with a CRISPR-GRIT gRNA targeting TRL1 (Guide #2) | This study |
| CaVip-7 | pV1093-CatRNA based integrative CRISPR-Cas9 system with a CRISPR-GRIT gRNA targeting TRL1 (Guide #3) | This study |
| CaVip-8 | pV1093-CatRNA based integrative CRISPR-Cas9 system with a CRISPR-GRIT gRNA targeting TRL1 (Guide #4) | This study |
| CaVip-9 | pV1093-CatRNA based integrative CRISPR-Cas9 system with a CRISPR- | This study |

| | | |
|---|---|---|
| | GRIT gRNA targeting RPB2 (Guide #1) | |
| CaVip-10 | pV1093-CatRNA based integrative CRISPR-Cas9 system with a CRISPR-GRIT gRNA targeting RPB2 (Guide #2) | This study |
| CaVip-11 | pV1093-CatRNA based integrative CRISPR-Cas9 system with a CRISPR-GRIT gRNA targeting RPB2 (Guide #3) | This study |
| CaVip-12 | pV1093-CatRNA based integrative CRISPR-Cas9 system with a CRISPR-GRIT gRNA targeting RPB2 (Guide #4) | This study |
| CaVip-13 | pV1093-CatRNA based integrative CRISPR-Cas9 system with a CRISPR-GRIT gRNA targeting RAD52 (Guide #1) | This study |
| CaVip-14 | pV1093-CatRNA based integrative CRISPR-Cas9 system with a CRISPR-GRIT gRNA targeting RAD52 (Guide #2) | This study |
| CaVip-15 | pV1093-CatRNA based integrative CRISPR-Cas9 system with a CRISPR-GRIT gRNA targeting RAD52 (Guide #3) | This study |
| CaVip-16 | pV1093-CatRNA based integrative CRISPR-Cas9 system with a CRISPR-GRIT gRNA targeting RAD52 (Guide #4) | This study |
| CaVip-17 | pV1093-CatRNA based integrative CRISPR-Cas9 system with a CRISPR-GRIT gRNA targeting RAD51 (Guide #1) | This study |
| CaVip-18 | pV1093-CatRNA based integrative CRISPR-Cas9 system with a CRISPR-GRIT gRNA targeting RAD51 (Guide #2) | This study |
| CaVip-19 | pV1093-CatRNA based integrative CRISPR-Cas9 system with a CRISPR-GRIT gRNA targeting RAD51 (Guide #3) | This study |
| CaVip-20 | pV1093-CatRNA based integrative CRISPR-Cas9 system with a CRISPR-GRIT gRNA targeting RAD51 (Guide #4) | This study |

| | | |
|---|---|---|
| CaVip-21 | pV1093-CatRNA based integrative CRISPR-Cas9 system with a CRISPR-GRIT gRNA targeting LIG4 (Guide #1) | This study |
| CaVip-22 | pV1093-CatRNA based integrative CRISPR-Cas9 system with a CRISPR-GRIT gRNA targeting LIG4 (Guide #2) | This study |
| CaVip-23 | pV1093-CatRNA based integrative CRISPR-Cas9 system with a CRISPR-GRIT gRNA targeting LIG4 (Guide #3) | This study |
| CaVip-24 | pV1093-CatRNA based integrative CRISPR-Cas9 system with a CRISPR-GRIT gRNA targeting LIG4 (Guide #4) | This study |
| CaVip3-NR | pV1093-CatRNA based integrative CRISPR-Cas9 system harboring gRNA CaVip3 (RPS15 gRNA #3) without a repair template | This study |
| CaVip7-NR | pV1093-CatRNA based integrative CRISPR-Cas9 system harboring gRNA CaVip7 (TRL1 gRNA #3) without a repair template | This study |
| CaVip12-NR | pV1093-CatRNA based integrative CRISPR-Cas9 system harboring gRNA CaVip12 (RPB2 gRNA #4) without a repair template | This study |
| CaVip15-NR | pV1093-CatRNA based integrative CRISPR-Cas9 system harboring gRNA CaVip15 (RAD52 gRNA #3) without a repair template | This study |
| CaVip17-NR | pV1093-CatRNA based integrative CRISPR-Cas9 system harboring gRNA CaVip17 (RAD51 gRNA #1) without a repair template | This study |
| CaVip23-NR | pV1093-CatRNA based integrative CRISPR-Cas9 system harboring gRNA CaVip23 (LIG4 gRNA #3) without a repair template | This study |
| CaVip 3 - 3 | pV1093-CatRNA based integrative CRISPR-Cas9 system harboring multiplexed CRISPR-GRIT gRNAs CaVip3 (RPS15 gRNA #3) with itself | This study |
| CaVip 3-Null | pV1093-CatRNA based integrative CRISPR-Cas9 system harboring | This study |

| | | |
|---|---|---|
| | multiplexed CRISPR-GRIT gRNAs CaVip3 (RPS15 gRNA #3) with a non-targeting gRNA | |
| CaVip 3 - 7 | pV1093-CatRNA based integrative CRISPR-Cas9 system harboring multiplexed CRISPR-GRIT gRNAs CaVip3 (RPS15 gRNA #3) and CaVip7 (TRL1 gRNA #3) | This study |
| CaVip 3 - 12 | pV1093-CatRNA based integrative CRISPR-Cas9 system harboring multiplexed CRISPR-GRIT gRNAs CaVip3 (RPS15 gRNA #3) and CaVip12 (RPB2 gRNA #4) | This study |
| CaVip 3 - 15 | pV1093-CatRNA based integrative CRISPR-Cas9 system harboring multiplexed CRISPR-GRIT gRNAs CaVip3 (RPS15 gRNA #3) and CaVip15 (RAD52 gRNA #3) | This study |
| CaVip 3 - 17 | pV1093-CatRNA based integrative CRISPR-Cas9 system harboring multiplexed CRISPR-GRIT gRNAs CaVip3 (RPS15 gRNA #3) and CaVip17 (RAD51 gRNA #1) | This study |
| CaVip 3 - 23 | pV1093-CatRNA based integrative CRISPR-Cas9 system harboring multiplexed CRISPR-GRIT gRNAs CaVip3 (RPS15 gRNA #3) and CaVip23 (LIG4 gRNA #3) | This study |
| CaVip 7 - 3 | pV1093-CatRNA based integrative CRISPR-Cas9 system harboring multiplexed CRISPR-GRIT gRNAs CaVip7 (TRL1 gRNA #3) and CaVip3 (RPS15 gRNA #3) | This study |
| CaVip 7 - 7 | pV1093-CatRNA based integrative CRISPR-Cas9 system harboring multiplexed CRISPR-GRIT gRNAs CaVip7 (TRL1 gRNA #3) with itself | This study |
| CaVip 7-Null | pV1093-CatRNA based integrative CRISPR-Cas9 system harboring multiplexed CRISPR-GRIT gRNAs CaVip7 (TRL1 gRNA #3) with a non-targeting gRNA | This study |
| CaVip 7 - 12 | pV1093-CatRNA based integrative CRISPR-Cas9 system harboring multiplexed CRISPR-GRIT gRNAs | This study |

| | | |
|---|---|---|
| | CaVip7 (TRL1 gRNA #3) and CaVip12 (RPB2 gRNA #4) | |
| CaVip 7 - 15 | pV1093-CatRNA based integrative CRISPR-Cas9 system harboring multiplexed CRISPR-GRIT gRNAs CaVip7 (TRL1 gRNA #3) and CaVip15 (RAD52 gRNA #3) | This study |
| CaVip 7 - 17 | pV1093-CatRNA based integrative CRISPR-Cas9 system harboring multiplexed CRISPR-GRIT gRNAs CaVip7 (TRL1 gRNA #3) and CaVip17 (RAD51 gRNA #1) | This study |
| CaVip 7 - 23 | pV1093-CatRNA based integrative CRISPR-Cas9 system harboring multiplexed CRISPR-GRIT gRNAs CaVip7 (TRL1 gRNA #3) and CaVip23 (LIG4 gRNA #3) | This study |
| CaVip 12 - 3 | pV1093-CatRNA based integrative CRISPR-Cas9 system harboring multiplexed CRISPR-GRIT gRNAs CaVip12 (RPB2 gRNA #4) and CaVip3 (RPS15 gRNA #3) | This study |
| CaVip 12 - 7 | pV1093-CatRNA based integrative CRISPR-Cas9 system harboring multiplexed CRISPR-GRIT gRNAs CaVip12 (RPB2 gRNA #4) and CaVip7 (TRL1 gRNA #3) | This study |
| CaVip 12 - 12 | pV1093-CatRNA based integrative CRISPR-Cas9 system harboring multiplexed CRISPR-GRIT gRNAs CaVip12 (RPB2 gRNA #4) with itself | This study |
| CaVip 12-Null | pV1093-CatRNA based integrative CRISPR-Cas9 system harboring multiplexed CRISPR-GRIT gRNAs CaVip12 (RPB2 gRNA #4) with a non-targeting gRNA | This study |
| CaVip 12 - 15 | pV1093-CatRNA based integrative CRISPR-Cas9 system harboring multiplexed CRISPR-GRIT gRNAs CaVip12 (RPB2 gRNA #4) and CaVip15 (RAD52 gRNA #3) | This study |
| CaVip 12 - 17 | pV1093-CatRNA based integrative CRISPR-Cas9 system harboring multiplexed CRISPR-GRIT gRNAs CaVip12 (RPB2 gRNA #4) and CaVip17 (RAD51 gRNA #1) | This study |

| | | |
|---|---|---|
| CaVip 12 - 23 | pV1093-CatRNA based integrative CRISPR-Cas9 system harboring multiplexed CRISPR-GRIT gRNAs CaVip12 (RPB2 gRNA #4) and CaVip23 (LIG4 gRNA #3) | This study |
| CaVip 15 - 3 | pV1093-CatRNA based integrative CRISPR-Cas9 system harboring multiplexed CRISPR-GRIT gRNAs CaVip15 (RAD52 gRNA #3) and CaVip3 (RPS15 gRNA #3) | This study |
| CaVip 15 - 7 | pV1093-CatRNA based integrative CRISPR-Cas9 system harboring multiplexed CRISPR-GRIT gRNAs CaVip15 (RAD52 gRNA #3) and CaVip7 (TRL1 gRNA #3) | This study |
| CaVip 15 - 12 | pV1093-CatRNA based integrative CRISPR-Cas9 system harboring multiplexed CRISPR-GRIT gRNAs CaVip15 (RAD52 gRNA #3) and CaVip12 (RPB2 gRNA #4) | This study |
| CaVip 15 - 17 | pV1093-CatRNA based integrative CRISPR-Cas9 system harboring multiplexed CRISPR-GRIT gRNAs CaVip15 (RAD52 gRNA #3) and CaVip17 (RAD51 gRNA #1) | This study |
| CaVip 15 - 23 | pV1093-CatRNA based integrative CRISPR-Cas9 system harboring multiplexed CRISPR-GRIT gRNAs CaVip15 (RAD52 gRNA #3) and CaVip23 (LIG4 gRNA #3) | This study |
| CaVip 17 - 3 | pV1093-CatRNA based integrative CRISPR-Cas9 system harboring multiplexed CRISPR-GRIT gRNAs CaVip17 (RAD51 gRNA #1) and CaVip3 (RPS15 gRNA #3) | This study |
| CaVip 17 - 7 | pV1093-CatRNA based integrative CRISPR-Cas9 system harboring multiplexed CRISPR-GRIT gRNAs CaVip17 (RAD51 gRNA #1) and CaVip7 (TRL1 gRNA #3) | This study |
| CaVip 17 - 12 | pV1093-CatRNA based integrative CRISPR-Cas9 system harboring multiplexed CRISPR-GRIT gRNAs CaVip17 (RAD51 gRNA #1) and CaVip12 (RPB2 gRNA #4) | This study |

| | | |
|---|---|---|
| CaVip 17 - 15 | pV1093-CatRNA based integrative CRISPR-Cas9 system harboring multiplexed CRISPR-GRIT gRNAs CaVip17 (RAD51 gRNA #1) and CaVip15 (RAD52 gRNA #3) | This study |
| CaVip 17 - 23 | pV1093-CatRNA based integrative CRISPR-Cas9 system harboring multiplexed CRISPR-GRIT gRNAs CaVip17 (RAD51 gRNA #1) and CaVip23 (LIG4 gRNA #3) | This study |
| CaVip 23 - 3 | pV1093-CatRNA based integrative CRISPR-Cas9 system harboring multiplexed CRISPR-GRIT gRNAs CaVip23 (LIG4 gRNA #3) and CaVip3 (RPS15 gRNA #3) | This study |
| CaVip 23 - 7 | pV1093-CatRNA based integrative CRISPR-Cas9 system harboring multiplexed CRISPR-GRIT gRNAs CaVip23 (LIG4 gRNA #3) and CaVip7 (TRL1 gRNA #3) | This study |
| CaVip 23 - 12 | pV1093-CatRNA based integrative CRISPR-Cas9 system harboring multiplexed CRISPR-GRIT gRNAs CaVip23 (LIG4 gRNA #3) and CaVip12 (RPB2 gRNA #4) | This study |
| CaVip 23 - 15 | pV1093-CatRNA based integrative CRISPR-Cas9 system harboring multiplexed CRISPR-GRIT gRNAs CaVip23 (LIG4 gRNA #3) and CaVip15 (RAD52 gRNA #3) | This study |
| CaVip 23 - 17 | pV1093-CatRNA based integrative CRISPR-Cas9 system harboring multiplexed CRISPR-GRIT gRNAs CaVip23 (LIG4 gRNA #3) and CaVip17 (RAD51 gRNA #1) | This study |
| CaVipNR 3-7 | pV1093-CatRNA based integrative CRISPR-Cas9 system harboring multiplexed gRNAs CaVip3 (RPS15 gRNA #3) and CaVip7 (TRL1 gRNA #3) without a repair template | This study |
| CaVipNR 3-12 | pV1093-CatRNA based integrative CRISPR-Cas9 system harboring multiplexed gRNAs CaVip3 (RPS15 gRNA #3) and CaVip12 (RPB2 gRNA #4) without a repair template | This study |

| | | |
|---|---|---|
| CaVipNR 3-15 | pV1093-CatRNA based integrative CRISPR-Cas9 system harboring multiplexed gRNAs CaVip3 (RPS15 gRNA #3) and CaVip15 (RAD52 gRNA #3) without a repair template | This study |
| CaVipNR 3-17 | pV1093-CatRNA based integrative CRISPR-Cas9 system harboring multiplexed gRNAs CaVip3 (RPS15 gRNA #3) and CaVip17 (RAD51 gRNA #1) without a repair template | This study |
| CaVipNR 3-23 | pV1093-CatRNA based integrative CRISPR-Cas9 system harboring multiplexed gRNAs CaVip3 (RPS15 gRNA #3) and CaVip23 (LIG4 gRNA #3) without a repair template | This study |
| pCC-CaCas9-ORI410 | pV1093-CatRNA based plasmid with the ORI410 sequence inserted in the NotI site upstream of the SNR52 promoter | This study |
| pCC-CaCas9-ORI1046 | pV1093-CatRNA based plasmid with the ORI1046 sequence inserted in the NotI site upstream of the SNR52 promoter | This study |
| pCC-CaCas9-ORI1046-GRIT ADE2 | pCC-CaCas9 based CRISPR-Cas9 system harboring an ADE2 targeting GRIT gRNA | This study |
| pCC-CaCas9-ORI1046-CaVip30 | pCC-CaCas9 based CRISPR-Cas9 system harboring an ACE2 targeting GRIT gRNA | This study |
| pCC-CaCas9-ORI1046-CaVip3 | pCC-CaCas9 based CRISPR-Cas9 system harboring gRNA CaVip3 | This study |
| pCC-CaCas9-ORI1046-CaVip7 | pCC-CaCas9 based CRISPR-Cas9 system harboring gRNA CaVip7 | This study |
| pCC-CaCas9-ORI1046-CaVip12 | pCC-CaCas9 based CRISPR-Cas9 system harboring gRNA CaVip12 | This study |
| pCC-CaCas9-ORI1046-CaVip15 | pCC-CaCas9 based CRISPR-Cas9 system harboring gRNA CaVip15 | This study |
| pCC-CaCas9-ORI1046-CaVip17 | pCC-CaCas9 based CRISPR-Cas9 system harboring gRNA CaVip17 | This study |
| pCC-CaCas9-ORI1046-CaVip23 | pCC-CaCas9 based CRISPR-Cas9 system harboring gRNA CaVip23 | This study |

**Table S3:** List of primers used in this study.

| Primer Name | Sequence (5'-3') | Description |
|---|---|---|
| CaVip-1-GRIT_FW | AAACGTCTCTGTCCATGGTTGGTCACTACTTGGGTGAATTCTCCATTACCTATACTCCAGTTAGAtaataaAGCTGGTAATGCTTCTTC | Forward primer for construction of CaVip-1 GRIT gRNA |
| CaVip-1-GRIT_RV | TTTCGTCTCCAAAACTGTCTAACTGGAGTATAGGTCACATCAAGTTTATCTCAATGGCATGAATTTAGAAGAAGCATTACCAGCTttat | Reverse primer for construction of CaVip-1 GRIT gRNA |
| CaVip-2-GRIT_FW | AAACGTCTCTGTCCATCAAAAAATTGAGAGCTGCCAGAGCTGCCACTGAACCAAATGAAAGACCAtaataaCAAAACTCACTTGAGAAA | Forward primer for construction of CaVip-2 GRIT gRNA |
| CaVip-2-GRIT_RV | TTTCGTCTCCAAAACTTGTCAAAACTCACTTGAGACAGAACCAATCATTTCTGGAACAACAATCATGTTTCTCAAGTGAGTTTTGttat | Reverse primer for construction of CaVip-2 GRIT gRNA |
| CaVip-3-GRIT_FW | AAACGTCTCTGTCCAGAAACATGATTGTTGTTCCAGAAATGATTGGTTCTGTTGTTGGTGTCTACtaataaAGTTTTCAACACTGTTGA | Forward primer for construction of CaVip-3 GRIT gRNA |
| CaVip-3-GRIT_RV | TTTCGTCTCCAAAACTTGTAGACACCAACAACAGACCAAGTAGTGACCAACCATTTCTGGTTTAATTTCAACAGTGTTGAAAACTttat | Reverse primer for construction of CaVip-3 GRIT gRNA |
| CaVip-4-GRIT_FW | AAACGTCTCTGTCCAGTAAAGTTTTCAACACTGTTGAAATTAAACCAGAAATGGTTGGTCACTACtaataaATTCTCCATTACCTATAC | Forward primer for construction of CaVip-4 GRIT gRNA |
| CaVip-4-GRIT_RV | TTTCGTCTCCAAAACAAGTAGTGACCAACCATTTCAAGCATTACCAGCTCTACCGTGTCTAACTGGAGTATAGGTAATGGAGAATttat | Reverse primer for construction of CaVip-4 GRIT gRNA |
| CaVip-5-GRIT_FW | AAACGTCTCTGTCCAAAAGTGAAATTGAACCAATTAATAAACAACTCCATATTACCATTGGTTGTtaataaAGCTACTGCTGTTGAGTC | Forward primer for construction of CaVip-5 GRIT gRNA |
| CaVip-5-GRIT_RV | TTTCGTCTCCAAAACGCCAGCTACTGCTGTTGAGTGATTGTCATACAATTCTTCCAATGTTATGTTTGACTCAACAGCAGTAGCTttat | Reverse primer for construction of CaVip-5 GRIT gRNA |
| CaVip-6-GRIT_FW | AAACGTCTCTGTCCAATGACGTTCTACAAAGTGAAATTGAACCAATTAATAAACA | Forward primer for construction of CaVip-6 GRIT gRNA |

| | | |
|---|---|---|
| | ACTCCATATTtaataaTTGTATCCCGCCAGCTAC | |
| CaVip-6-GRIT_RV | TTTCGTCTCCAAAACTTGGTTGTATCCCGCCAGCTATTCTTCCAATGTTATGTTTGACTCAACAGCAGTAGCTGGCGGGATACAAttat | Reverse primer for construction of CaVip-6 GRIT gRNA |
| CaVip-7-GRIT_FW | AAACGTCTCTGTCCACTCAGATCAAATTAGATTCTTGGAAATTTTTGGAATGGGATTATGGCAAAtaataaTCAACTTCCAATTCAAGC | Forward primer for construction of CaVip-7 GRIT gRNA |
| CaVip-7-GRIT_RV | TTTCGTCTCCAAAACAAGGTTTGCCATAATCCCATTAGTATCATTATTTAATGTAAACAATCCACGCGCTTGAATTGGAAGTTGAttat | Reverse primer for construction of CaVip-7 GRIT gRNA |
| CaVip-8-GRIT_FW | AAACGTCTCTGTCCAATGGTCGAGAAATTGAAGGGTTTGTGATAAGATGTCACCGCCAACTGCATtaataaTGACACTGACGGGGATTG | Forward primer for construction of CaVip-8 GRIT gRNA |
| CaVip-8-GRIT_RV | TTTCGTCTCCAAAACATGGTGACACTGACGGGGATATGGCTGTTCAAATTTATACTTGAAAAAAAAGCAATCCCCGTCAGTGTCAttat | Reverse primer for construction of CaVip-8 GRIT gRNA |
| CaVip-9-GRIT_FW | AAACGTCTCTGTCCAGAATTAAGCCAACCACCAGTGGCAATACTCATACATACACACATTGTGAAtaataaGTCTATGATATTGGGTGT | Forward primer for construction of CaVip-9 GRIT gRNA |
| CaVip-9-GRIT_RV | TTTCGTCTCCAAAACTCCGTCTATGATATTGGGTGTATGATCTGGGAACGGAATAATTGAGGCGGCAACACCCAATATCATAGACttat | Reverse primer for construction of CaVip-9 GRIT gRNA |
| CaVip-10-GRIT_FW | AAACGTCTCTGTCCAATGGTGGAGACTTTAATGTTACTTTGGCTGTTAAATCACAGACAATCACCtaataaAAGATACTCGTTGGCTAC | Forward primer for construction of CaVip-10 GRIT gRNA |
| CaVip-10-GRIT_RV | TTTCGTCTCCAAAACTCGGTGATTGTCTGTGATTTTCATGGCTTTTCTTTGCTCACCCCAATTACCAGTAGCCAACGAGTATCTTttat | Reverse primer for construction of CaVip-10 GRIT gRNA |
| CaVip-11-GRIT_FW | AAACGTCTCTGTCCAGATTAAAACACGGTACATATGAGAAATTAGATGAAGATGGGTTGATCGCCtaataaCAGAGTTAGTGGTGAAGA | Forward primer for construction of CaVip-11 GRIT gRNA |

| | | |
|---|---|---|
| CaVip-11-GRIT_RV | TTTCGTCTCCAAAACCAGGGGCGATCAACCCATCTCAGGTATAGGAGTTGTTTTACCAATAATAATATCTTCACCACTAACTCTGttat | Reverse primer for construction of CaVip-11 GRIT gRNA |
| CaVip-12-GRIT_FW | AAACGTCTCTGTCCAGATTAAAACACGGTACATATGAGAAATTAGATGAAGATGGGTTGATCGCCtaataaCAGAGTTAGTGGTGAAGA | Forward primer for construction of CaVip-12 GRIT gRNA |
| CaVip-12-GRIT_RV | TTTCGTCTCCAAAACAGGGGCGATCAACCCATCTTCAGGTATAGGAGTTGTTTTACCAATAATAATATCTTCACCACTAACTCTGttat | Reverse primer for construction of CaVip-12 GRIT gRNA |
| CaVip-13-GRIT_FW | AAACGTCTCTGTCCACAATTAAACGAAGTGATATTAAAAATAGTGTCGCTACGACACCGTCACCAtaataaAAACACATCTAGTAATCG | Forward primer for construction of CaVip-13 GRIT gRNA |
| CaVip-13-GRIT_RV | TTTCGTCTCCAAAACAATGGTGACGGTGTCGTAGCTATTGTTTGATGTTATCATGGGTTTATTAATTCGATTACTAGATGTGTTTttat | Reverse primer for construction of CaVip-13 GRIT gRNA |
| CaVip-14-GRIT_FW | AAACGTCTCTGTCCAAAGAACAAAAGAGACAACAAGAAGACGCGTCAAGGTTGGATAATTCAGGTtaataaGCAGCACCAGCAACAAAA | Forward primer for construction of CaVip-14 GRIT gRNA |
| CaVip-14-GRIT_RV | TTTCGTCTCCAAAACGTTGACCTGAATTATCCAACGAATTGATCCGTTTGAAACTACGTTTGAGTTATTTTGTTGCTGGTGCTGCttat | Reverse primer for construction of CaVip-14 GRIT gRNA |
| CaVip-15-GRIT_FW | AAACGTCTCTGTCCAGACCACCACAACAACAACCTCAGCAACCTCAGCAACCTCAACCCAACCAAtaataaTCCCTTTCGACCAGACGA | Forward primer for construction of CaVip-15 GRIT gRNA |
| CaVip-15-GRIT_RV | TTTCGTCTCCAAAACCTGTTGGTTGGGTTGAGGTTCAAGTGGTTGATTCAAACGACGAGCTCGTGACTCGTCTGGTCGAAAGGGAttat | Reverse primer for construction of CaVip-15 GRIT gRNA |
| CaVip-16-GRIT_FW | AAACGTCTCTGTCCACTCCGCAACCACGACCACCACAACAACAACCTCAGCAACCTCAGCAACCTtaataaCCAACAGAGGCTTCCCTT | Forward primer for construction of CaVip-16 GRIT gRNA |
| CaVip-16-GRIT_RV | TTTCGTCTCCAAAACAACCAACAGAGGCTTCCCTTTCAAACGACGAGCTCGTGACTCGTCTGGTCGAAAGGGAAGCCTCTGTTGGttat | Reverse primer for construction of CaVip-16 GRIT gRNA |

| | | |
|---|---|---|
| CaVip-17-GRIT_FW | AAACGTCTCTGTCCAAAGTTGATGGTATGTCTGGTATGTTTAATCCTGATCCTAAGAAACCAATTtaataaCATTATTGCCCATTCATC | Forward primer for construction of CaVip-17 GRIT gRNA |
| CaVip-17-GRIT_RV | TTTCGTCTCCAAAACCCAATTGGTTTCTTAGGATCCTCTTCCCTTTTTAAACGATAATCTAGTTGTTGATGAATGGGCAATAATGttat | Reverse primer for construction of CaVip-17 GRIT gRNA |
| CaVip-18-GRIT_FW | AAACGTCTCTGTCCACTCAAGTTGATGGTATGTCTGGTATGTTTAATCCTGATCCTAAGAAACCAtaataaTAACATTATTGCCCATTC | Forward primer for construction of CaVip-18 GRIT gRNA |
| CaVip-18-GRIT_RV | TTTCGTCTCCAAAACCAATTGGTTTCTTAGGATCATTCCCTTTTTAAACGATAATCTAGTTGTTGATGAATGGGCAATAATGTTAttat | Reverse primer for construction of CaVip-18 GRIT gRNA |
| CaVip-19-GRIT_FW | AAACGTCTCTGTCCATTGTTGCTCAAGTTGATGGTATGTCTGGTATGTTTAATCCTGATCCTAAGtaataaTGGGGGTAACATTATTGC | Forward primer for construction of CaVip-19 GRIT gRNA |
| CaVip-19-GRIT_RV | TTTCGTCTCCAAAACATTGGGGGTAACATTATTGCTTTTAAACGATAATCTAGTTGTTGATGAATGGGCAATAATGTTACCCCCAttat | Reverse primer for construction of CaVip-19 GRIT gRNA |
| CaVip-20-GRIT_FW | AAACGTCTCTGTCCATTAGAACAGGTAAATCACAATTATGTCACACGTTAGCCGTTACGTGTCAGtaataaTGATATGGGTGGTGGTGA | Forward primer for construction of CaVip-20 GRIT gRNA |
| CaVip-20-GRIT_RV | TTTCGTCTCCAAAACATTGATATGGGTGGTGGTGATACCTTCAGTATCAATGTAAAGACATTTTCCTTCACCACCACCCATATCAttat | Reverse primer for construction of CaVip-20 GRIT gRNA |
| CaVip-21-GRIT_FW | AAACGTCTCTGTCCAATGACTTACAAAAGACATTACAATTTTCAACAAATCCCGATTTGAGACTAtaataaGCAGTTAGCTATACATCC | Forward primer for construction of CaVip-21 GRIT gRNA |
| CaVip-21-GRIT_RV | TTTCGTCTCCAAAACTCTGCAGTTAGCTATACATCTTTCTGACAACTGTGGTTTAAACTTGAAACAAGGATGTATAGCTAACTGCttat | Reverse primer for construction of CaVip-21 GRIT gRNA |
| CaVip-22-GRIT_FW | AAACGTCTCTGTCCAATAAAAATGGTGGGTCTTCGAGAATTCCACATTTTGTTACTGAGGCATTCtaataaTTCAATCAAGATGAATTA | Forward primer for construction of CaVip-22 GRIT gRNA |

| | | |
|---|---|---|
| CaVip-22-GRIT_RV | TTTCGTCTCCAAAACGAACGAATGCCTCAGTAACAACCTAAACTTGTAATCGTCAGGATCGGGAATATAATTCATCTTGATTGAAttat | Reverse primer for construction of CaVip-22 GRIT gRNA |
| CaVip-23-GRIT_FW | AAACGTCTCTGTCCAAAAGTGAACTTGAAGACCTTCGTAATGGATTTGATTGGGGTGATCTTAAAtaataaCTACTTGTTCAAAGGTTT | Forward primer for construction of CaVip-23 GRIT gRNA |
| CaVip-23-GRIT_RV | TTTCGTCTCCAAAACAGGTTTAAGATCACCCCAATCACTCAAGTTATTCCCACACACGTAAAATGACAAACCTTTGAACAAGTAGttat | Reverse primer for construction of CaVip-23 GRIT gRNA |
| CaVip-24-GRIT_FW | AAACGTCTCTGTCCATATTGAAGAATGTACAGCTGAAGTATGAAATTGATGGGTTCAGAAACCCAtaataaAAAAGTCAAACCAGAGTA | Forward primer for construction of CaVip-24 GRIT gRNA |
| CaVip-24-GRIT_RV | TTTCGTCTCCAAAACATCTGGGTTTCTGAACCCATCAAGATCTAAATTTTCACCAAATTTTTCTAAATACTCTGGTTTGACTTTTttat | Reverse primer for construction of CaVip-24 GRIT gRNA |
| CaVip-30-GRIT_FW | AAACGTCTCTGTCCATGCATTTATCACCTTTGAAAAAACAATTACCAAACACTCCCACAAAGCAAtaataaCACCATTGAATGGAGTCC | Forward primer for construction of CaVip-30 (*ACE2*) GRIT gRNA for insertion into pCC-CaCas9-ORI1046 |
| CaVip-30-GRIT_RV | TTTCGTCTCCAAAACCTGTCACCATTGAATGGAGTGTAATGGTTGCTTTGAGTTTGGTGATATAACTGGACTCCATTCAATGGTGttat | Reverse primer for construction of CaVip-30 (*ACE2*) GRIT gRNA for insertion into pCC-CaCas9-ORI1046 |
| CC-373 | AAACGTCTCTGTCCAGAATCAACCCCATCTAATGTAGATCCCTTAACTGGAACACCAATGACAGGCTATTACATGGCCGCCACCATACCT | Forward primer for the construction of the ADE2 GRIT gRNA for insertion into pCC-CaCas9-ORI1046 |
| HiCRISPRRV | TTTCGTCTCCAAAACCATTGCCTGTCATTGGTGTTTTGCTGGTGCAGGTGGTGCTGCTCATTTGCCAGGTATGGTGGCGGCCATG | Reverse primer for the construction of the ADE2 GRIT gRNA for insertion into pCC-CaCas9-ORI1046 |
| CC-518 | gtccaTCTGTTGTTGGTGTCTACAAg | Forward oligo for the CaVip3 (RPS15) gRNA without a repair template |
| CC-519 | aaaacTTGTAGACACCAACAACAGAt | Reverse oligo for the CaVip3 (RPS15) gRNA without a repair template |

| | | |
|---|---|---|
| CC-520 | gtccaATGGGATTATGGCAAACCTTg | Forward oligo for the CaVip7 (TRL1) gRNA without a repair template |
| CC-521 | aaaacAAGGTTTGCCATAATCCCATt | Reverse oligo for the CaVip7 (TRL1) gRNA without a repair template |
| CC-522 | gtccaAAGATGGGTTGATCGCCCCTg | Forward oligo for the CaVip12 (RPB2) gRNA without a repair template |
| CC-523 | aaaacAGGGGCGATCAACCCATCTTt | Reverse oligo for the CaVip12 (RPB2) gRNA without a repair template |
| CC-524 | gtccaAACCTCAACCCAACCAACAGg | Forward oligo for the CaVip15 (RAD52) gRNA without a repair template |
| CC-525 | aaaacCTGTTGGTTGGGTTGAGGTTt | Reverse oligo for the CaVip15 (RAD52) gRNA without a repair template |
| CC-526 | gtccaGATCCTAAGAAACCAATTGGg | Forward oligo for the CaVip17 (RAD51) gRNA without a repair template |
| CC-527 | aaaacCCAATTGGTTTCTTAGGATCt | Reverse oligo for the CaVip17 (RAD51) gRNA without a repair template |
| CC-528 | gtccaATTGGGGTGATCTTAAACCTg | Forward oligo for the CaVip23 (LIG4) gRNA without a repair template |
| CC-529 | aaaacAGGTTTAAGATCACCCCAATt | Reverse oligo for the CaVip23 (LIG4) gRNA without a repair template |
| CC-932 | gaaaagatcgttttctttattattctctagttttgacggcGGAACATCTGAAATTGGTTC | ORI 410 Forward primer with Gibson overhangs for insertion into the pV1093-CatRNA NotI site |
| CC-933 | tgtgcttgatttgaacggactaagtctaatcactgcggccTTGATGATTGGATCGGGTTC | ORI 410 Reverse primer with Gibson overhangs for insertion into the pV1093-CatRNA NotI site |
| CC-934 | gaaaagatcgttttctttattattctctagttttgacggcATATATTTGTGATTCAACCACAC | ORI 1046 Forward primer with Gibson overhangs for insertion into the pV1093-CatRNA NotI site |
| CC-935 | tgtgcttgatttgaacggactaagtctaatcactgcggccCAAAAATATCTCGTGAATCTTTTC | ORI 1046 Reverse primer with Gibson overhangs for insertion into the pV1093-CatRNA NotI site |

| | | |
|---|---|---|
| CC-936 | CCGTACACCAAGAAGTTAGACATCCGTACACCAAcaccagtctttgagaaattctcaaaacc | Foward primer for linearization and 1.5x telomere repeat addition to linear plasmid based CRISPR-Cas systems |
| CC-937 | CCGTACACCAAGAAGTTAGACATCCGTACACCAAgacgttagcatcgattaataaaatgtgc | Reverse primer for linearization and 1.5x telomere repeat addition to linear plasmid based CRISPR-Cas systems |
| CC-938 | caccagtctttgagaaattctcaaaacc | Foward primer for linearization of linear plasmid based CRISPR-Cas systems (No telomere repeats) |
| CC-939 | gacgttagcatcgattaataaaatgtgc | Reverse primer for linearization of linear plasmid based CRISPR-Cas systems (No telomere repeats) |
| CC-21 | ctcgacaacaaaaaagtttgatctg | Foward primer for checking ORI sequence insertion |
| CC-22 | gttggtggggcaataactcc | Reverse primer for checking gRNA and ORI sequence insertions |
| CC-79 | gcaccaattacgtaccaag | Foward primer for checking and sequencing gRNA insertions |
| CC-377 | AAACGTCTCTATTTGCAAAACGGGCGTGTGG | Multiplexing gRNA #1 FW |
| CC-380 | TTTCGTCTCAAAAAAgcaccgactcggtgc | Multiplexing gRNA #2 RV |
| CC-463 | TTTCGTCTCgcgtagtcatgagtcagacttatcattatccttaaacactcg | Discrete Transcriptional Unit Multiplex gRNA #1 RV primer |
| CC-464 | AAACGTCTCTTACGgtgattagacttagtccgttc | Discrete Transcriptional Unit Multiplex gRNA #2 FW primer |

**Table S4:** List of gRNAs and CRISPR-GRIT sequences used in this study.

| **Plasmid** | **gRNA Seed Sequence (5'-3')** | **CRISPR-GRIT Sequence (Repair Template + Seed)** |
|---|---|---|
| CaVip-1 | ACCTATACTCCAGTTAGACA | TGGTTGGTCACTACTTGGGTGAATTCTCCATTACCTATACTCCAGTTAGAtaataaAGCTGGTAATGCTTCTTCTAAATTCATGCCATTGAGATAAACTTGATGTGACCTATACTCCAGTTAGACA |
| CaVip-2 | TCTCAAGTGAGTTTTGACAA | TCAAAAAATTGAGAGCTGCCAGAGCTGCCACTGAACCAAATGAAAGACCAtaataaCAAAACTCACTTGAGAAACATGATTGTTGTTCCAGAAATGATTGGTTCTGTCTCAAGTGAGTTTTGACAA |
| CaVip-3 | TCTGTTGTTGGTGTCTACAA | GAAACATGATTGTTGTTCCAGAAATGATTGGTTCTGTTGTTGGTGTCTACtaataaAGTTTTCAACACTGTTGAAATTAAACCAGAAATGGTTGGTCACTACTTGGTCTGTTGTTGGTGTCTACAA |
| CaVip-4 | GAAATGGTTGGTCACTACTT | GTAAAGTTTTCAACACTGTTGAAATTAAACCAGAAATGGTTGGTCACTACtaataaATTCTCCATTACCTATACTCCAGTTAGACACGGTAGAGCTGGTAATGCTTGAAATGGTTGGTCACTACTT |
| CaVip-5 | ACTCAACAGCAGTAGCTGGC | AAAGTGAAATTGAACCAATTAATAAACAACTCCATATTACCATTGGTTGTtaataaAGCTACTGCTGTTGAGTCAAACATAACATTGGAAGAATTGTATGACAATCACTCAACAGCAGTAGCTGGC |
| CaVip-6 | AGCTGGCGGGATACAACCAA | ATGACGTTCTACAAAGTGAAATTGAACCAATTAATAAACAACTCCATATTtaataaTTGTATCCCGCCAGCTACTGCTGTTGAGTCAAACATAACATTGGAAGAATAGCTGGCGGGATACAACCAA |
| CaVip-7 | ATGGGATTATGGCAAACCTT | CTCAGATCAAATTAGATTCTTGGAAATTTTTGGAATGGGATTATGGCAAAtaataaTCAACTTCCAATTCAAGCGCGTGGATTGTTTACATTAAATAATGATACTAATGGGATTATGGCAAACCTT |

| | | |
|---|---|---|
| CaVip-8 | ATCCCCGTCAGTGTCACCAT | ATGGTCGAGAAATTGAAGGGTTT<br>GTGATAAGATGTCACCGCCAACT<br>GCATtaataaTGACACTGACGGGGA<br>TTGCTTTTTTTTCAAGTATAAATT<br>TGAACAGCCATATCCCCGTCAGT<br>GTCACCAT |
| CaVip-9 | CACCCAATATCATAGACGGA | GAATTAAGCCAACCACCAGTGGC<br>AATACTCATACATACACACATTG<br>TGAAtaataaGTCTATGATATTGGGT<br>GTTGCCGCCTCAATTATTCCGTTC<br>CCAGATCATACACCCAATATCAT<br>AGACGGA |
| CaVip-10 | AAATCACAGACAATCACCGA | ATGGTGGAGACTTTAATGTTACT<br>TTGGCTGTTAAATCACAGACAAT<br>CACCtaataaAAGATACTCGTTGGCT<br>ACTGGTAATTGGGGTGAGCAAAG<br>AAAAGCCATGAAAATCACAGAC<br>AATCACCGA |
| CaVip-11 | AGATGGGTTGATCGCCCCTG | GATTAAAACACGGTACATATGAG<br>AAATTAGATGAAGATGGGTTGAT<br>CGCCtaataaCAGAGTTAGTGGTGA<br>AGATATTATTATTGGTAAAACAA<br>CTCCTATACCTGAGATGGGTTGA<br>TCGCCCCTG |
| CaVip-12 | AAGATGGGTTGATCGCCCCT | GATTAAAACACGGTACATATGAG<br>AAATTAGATGAAGATGGGTTGAT<br>CGCCtaataaCAGAGTTAGTGGTGA<br>AGATATTATTATTGGTAAAACAA<br>CTCCTATACCTGAAGATGGGTTG<br>ATCGCCCCT |
| CaVip-13 | GCTACGACACCGTCACCATT | CAATTAAACGAAGTGATATTAAA<br>AATAGTGTCGCTACGACACCGTC<br>ACCAtaataaAAACACATCTAGTAA<br>TCGAATTAATAAACCCATGATAA<br>CATCAAACAATAGCTACGACACC<br>GTCACCATT |
| CaVip-14 | GTTGGATAATTCAGGTCAAC | AAGAACAAAAGAGACAACAAGA<br>AGACGCGTCAAGGTTGGATAATT<br>CAGGTtaataaGCAGCACCAGCAAC<br>AAAATAACTCAAACGTAGTTTCA<br>AACGGATCAATTCGTTGGATAAT<br>TCAGGTCAAC |
| CaVip-15 | AACCTCAACCCAACCAACAG | GACCACCACAACAACAACCTCAG<br>CAACCTCAGCAACCTCAACCCAA<br>CCAAtaataaTCCCTTTCGACCAGAC<br>GAGTCACGAGCTCGTCGTTTGAA |

| | | |
|---|---|---|
| | | TCAACCACTTGAACCTCAACCCA<br>ACCAACAG |
| CaVip-16 | AAGGGAAGCCTCTGTTGGTT | CTCCGCAACCACGACCACCACAA<br>CAACAACCTCAGCAACCTCAGCA<br>ACCTtaataaCCAACAGAGGCTTCCC<br>TTTCGACCAGACGAGTCACGAGC<br>TCGTCGTTTGAAAGGGAAGCCTC<br>TGTTGGTT |
| CaVip-17 | GATCCTAAGAAACCAATTGG | AAGTTGATGGTATGTCTGGTATG<br>TTTAATCCTGATCCTAAGAAACC<br>AATTtaataaCATTATTGCCCATTCA<br>TCAACAACTAGATTATCGTTTAA<br>AAAGGGAAGAGGATCCTAAGAA<br>ACCAATTGG |
| CaVip-18 | TGATCCTAAGAAACCAATTG | CTCAAGTTGATGGTATGTCTGGT<br>ATGTTTAATCCTGATCCTAAGAA<br>ACCAtaataaTAACATTATTGCCCAT<br>TCATCAACAACTAGATTATCGTT<br>TAAAAAGGGAATGATCCTAAGA<br>AACCAATTG |
| CaVip-19 | GCAATAATGTTACCCCCAAT | TTGTTGCTCAAGTTGATGGTATG<br>TCTGGTATGTTTAATCCTGATCCT<br>AAGtaataaTGGGGGTAACATTATT<br>GCCCATTCATCAACAACTAGATT<br>ATCGTTTAAAAGCAATAATGTTA<br>CCCCCAAT |
| CaVip-20 | TCACCACCACCCATATCAAT | TTAGAACAGGTAAATCACAATTA<br>TGTCACACGTTAGCCGTTACGTG<br>TCAGtaataaTGATATGGGTGGTGGT<br>GAAGGAAAATGTCTTTACATTGA<br>TACTGAAGGTATCACCACCACCC<br>ATATCAAT |
| CaVip-21 | GATGTATAGCTAACTGCAGA | ATGACTTACAAAAGACATTACAA<br>TTTTCAACAAATCCCGATTTGAG<br>ACTAtaataaGCAGTTAGCTATACAT<br>CCTTGTTTCAAGTTTAAACCACA<br>GTTGTCAGAAAGATGTATAGCTA<br>ACTGCAGA |
| CaVip-22 | TGTTACTGAGGCATTCGTTC | ATAAAAATGGTGGGTCTTCGAGA<br>ATTCCACATTTTGTTACTGAGGC<br>ATTCtaataaTTCAATCAAGATGAAT<br>TATATTCCCGATCCTGACGATTA<br>CAAGTTTAGGTTGTTACTGAGGC<br>ATTCGTTC |

| | | |
|---|---|---|
| CaVip-23 | ATTGGGGTGATCTTAAACCT | AAAGTGAACTTGAAGACCTTCGTAATGGATTTGATTGGGGTGATCTTAAAtaataaCTACTTGTTCAAAGGTTTGTCATTTTACGTGTGTGGGAATAACTTGAGTGATTGGGGTGATCTTAAACCT |
| CaVip-24 | ATGGGTTCAGAAACCCAGAT | TATTGAAGAATGTACAGCTGAAGTATGAAATTGATGGGTTCAGAAACCCAtaataaAAAAGTCAAACCAGAGTATTTAGAAAAATTTGGTGAAAATTTAGATCTTGATGGGTTCAGAAACCCAGAT |
| CaVip-ADE2 | AACACCAATGACAGGCAATG | GAATCAACCCCATCTAATGTAGATCCCTTAACTGGAACACCAATGACAGGctattaCATGGCCGCCACCATACCTGGCAAATGAGCAGCACCACCTGCACCAGCAAAACACCAATGACAGGCAATG |
| CaVip-30 | TCTGTTGGTTGAGGCTGCTT | CATCTGGACAAACAACACCAAACATGTCACAACCTCCTAGTGCTGGCACGtaataaGCAGCCTCAACCAACAGAGCAAATGCGTCAATTACAAGACAAGCAACAGCTCTGTTGGTTGAGGCTGCTT |

**Table S5.** List of single target gRNAs with on-target scores and $\log_2$ colony reductions (Log2CR) using the CRISPR-GRIT designs.

| Plasmid | Gene Target | gRNA Seed Sequence (5'-3') | CASPER On-Target Score | CRISPR-GRIT Log2CR (Average ± SD) |
|---|---|---|---|---|
| CaVip-1 | *RPS15* | ACCTATACTCCAGTTAGACA | 20 | 2.94 ± 0.22 |
| CaVip-2 | *RPS15* | TCTCAAGTGAGTTTTGACAA | 54 | 1.19 ± 0.36 |
| CaVip-3 | *RPS15* | TCTGTTGTTGGTGTCTACAA | 74 | 3.99 ± 0.57 |
| CaVip-4 | *RPS15* | GAAATGGTTGGTCACTACTT | 65 | 1.55 ± 0.55 |
| CaVip-5 | *TRL1* | ACTCAACAGCAGTAGCTGGC | 46 | -0.20 ± 0.27 |
| CaVip-6 | *TRL1* | AGCTGGCGGGATACAACCAA | 94 | 0.83 ± 0.34 |
| CaVip-7 | *TRL1* | ATGGGATTATGGCAAACCTT | 100 | 2.60 ± 0.33 |
| CaVip-8 | *TRL1* | ATCCCCGTCAGTGTCACCAT | 100 | 0.38 ± 0.47 |
| CaVip-9 | *RPB2* | CACCCAATATCATAGACGGA | 58 | 0.07 ± 0.38 |
| CaVip-10 | *RPB2* | AAATCACAGACAATCACCGA | 37 | 0.22 ± 0.45 |
| CaVip-11 | *RPB2* | AGATGGGTTGATCGCCCCTG | 97 | 0.63 ± 0.41 |
| CaVip-12 | *RPB2* | AAGATGGGTTGATCGCCCCT | 91 | 2.53 ± 0.63 |
| CaVip-13 | *RAD52* | GCTACGACACCGTCACCATT | 40 | 0.43 ± 0.88 |
| CaVip-14 | *RAD52* | GTTGGATAATTCAGGTCAAC | 38 | 0.03 ± 0.48 |
| CaVip-15 | *RAD52* | AACCTCAACCCAACCAACAG | 100 | 3.01 ± 0.3 |
| CaVip-16 | *RAD52* | AAGGGAAGCCTCTGTTGGTT | 100 | 0.61 ± 0.33 |
| CaVip-17 | *RAD51* | GATCCTAAGAAACCAATTGG | 52 | 1.21 ± 0.48 |
| CaVip-18 | *RAD51* | TGATCCTAAGAAACCAATTG | 40 | 0.25 ± 0.42 |
| CaVip-19 | *RAD51* | GCAATAATGTTACCCCCAAT | 100 | 0.80 ± 0.5 |
| CaVip-20 | *RAD51* | TCACCACCACCCATATCAAT | 100 | 0.63 ± 0.11 |
| CaVip-21 | *LIG4* | GATGTATAGCTAACTGCAGA | 38 | -0.03 ± 0.36 |
| CaVip-22 | *LIG4* | TGTTACTGAGGCATTCGTTC | 39 | -0.21 ± 0.15 |
| CaVip-23 | *LIG4* | ATTGGGGTGATCTTAAACCT | 84 | 1.04 ± 0.39 |
| CaVip-24 | *LIG4* | ATGGGTTCAGAAACCCAGAT | 83 | -0.29 ± 0.19 |
